# A functional corona around extracellular vesicles enhances angiogenesis during skin regeneration and signals in immune cells

**DOI:** 10.1101/808808

**Authors:** Martin Wolf, Rodolphe W Poupardin, Patricia Ebner-Peking, André Cronemberger Andrade, Constantin Blöchl, Astrid Obermayer, Fausto Gueths Gomes, Balazs Vari, Essi Eminger, Heide-Marie Binder, Anna M Raninger, Sarah Hochmann, Gabriele Brachtl, Andreas Spittler, Thomas Heuser, Racheli Ofir, Christian G Huber, Zami Aberman, Katharina Schallmoser, Hans-Dieter Volk, Dirk Strunk

## Abstract

Nanoparticles can acquire a protein corona defining their biological identity. Corona functions were not yet considered for cell-derived extracellular vesicles (EVs). Here we demonstrate that nanosized EVs from therapy-grade human placental-expanded (PLX) stromal cells are surrounded by an imageable and functional protein corona when enriched with permissive technology. Scalable EV separation from cell-secreted soluble factors via tangential flow-filtration and subtractive tandem mass-tag proteomics revealed significant enrichment of predominantly immunomodulatory and proangiogenic proteins. Western blot, calcein-based flow cytometry, super-resolution and electron microscopy verified EV identity. PLX-EVs protected corona proteins from protease digestion. EVs significantly ameliorated human skin regeneration and angiogenesis in vivo, induced differential signaling in immune cells, and dose-dependently inhibited T cell proliferation in vitro. Corona removal by size-exclusion or ultracentrifugation abrogated angiogenesis. Re-establishing an artificial corona by cloaking EVs with defined proangiogenic proteins served as a proof-of-concept. Understanding EV corona formation will improve rational EV-inspired nanotherapy design.

---

EVs are a heterogeneous family of generally nanosized membrane-coated vesicular structures derived from virtually all cell types^1^. EVs comprise prototypic endosomal-derived exosomes, assembled and released via multivesicular bodies, as well as outer cell membrane-derived sub-micron sized ectosomes (microvesicles) and apoptotic bodies^2, 3^. Conceptually, EVs transport their cargo to target sites enabling action over distance^3, 4^. Address codes on the EV surface may contribute to target specificity^5^. In contrast, the biological identity of synthetic nanoparticles is critically defined by their corona acquired upon entry into protein-rich environments. Albumin, representing the most abundant plasma protein, can act as a dysopsonin inhibiting nanoparticle uptake^6^. Other distinct corona components can further decrease or increase cellular nanoparticle uptake in a biological environment^7^. It is meanwhile well established that rapid plasma protein corona formation, within seconds, can determine nanoparticle function^8^. Vice versa, nanoparticles themselves can protect protein conjugates in their corona from protease degradation, thus also shaping functionality^9^. This complex interplay impacts spatial and temporal distribution of nanotherapeutics and may, at least in part, contribute to clinical failure of certain targeted nanomedicines^10^. The impact of a corona formation phenomenon on EV biology and function has not been addressed so far.

Peripheral artery disease (PAD) affects more than 15% of the >80 year-old population worldwide^11^. Critical limb ischemia (CLI) is an end-stage of PAD resulting in high amputation rates and is associated with increased risk for cardiovascular events and death. Allogeneic PLX stromal cells are currently evaluated in a clinical phase III trial (<u>NCT03006770</u>) for efficiency as an advanced CLI therapy^12^. The regenerative potential of placental cells is not restricted to CLI as evidenced by their hematopoietic support activity^13^, capacity to protect from radiation injury^14^, their current investigation for improving muscle regeneration in patients after hip arthroplasty^15^ and for treatment of coronavirus disease 2019 (COVID-19) complications. Their mode of action was related to cytokine and growth factor secretion promoting angiogenesis, cell recruitment, migration and proliferation, resulting in tissue regeneration^16^. The immunomodulatory capacity of stromal cells has also been hypothesized to exert beneficial effects on local and systemic immune responses^17^.

This study was inspired by the multiplicity of EV functions during intercellular communication, paving their way towards clinical applicability, and partly replacing conventional cell-based and nanotherapies^18, 19^. We hypothesized that PLX-EVs contribute to the mode of action of cell-based therapy. We used conditioned media (CM) obtained after short-term propagation of clinical grade PLX cell products^12^ under animal serum-free particle-depleted conditions to separate EVs from corresponding PLX-derived soluble factors for comparative proteomic and functional analysis (**Fig.S1**). Initial results confirmed proangiogenic and immunomodulatory potency of PLX-EVs. Surprisingly, further EV purification by size exclusion chromatography (SEC) or ultracentrifugation abrogated EV function, but could be re-established by cloaking PLX-EVs with a protein corona comprising three selected proangiogenic proteins dissolved in human albumin solution.

## RESULTS

### Physically defined media with a low particle count permit cell-derived EV analysis

In initial experiments, we determined the particle content of standard media for stromal cell culture, because high preexisting particle concentrations would disable PLX-EV purification. In fact, alpha-modified minimum essential medium (α-MEM) and other conventional media supplemented with either fetal bovine serum (FBS) or pooled human platelet lysate (HPL)^20^ contained mean 4 x 10^8^ – 3 x 10^9^ particles/mL. Fibrinogen-depleted α-MEM (α-MEM*), which could be used without heparin that otherwise might inhibit EV uptake^21, 22^, had even more particles of up to 10^10^/mL. Both ultracentrifugation (UCF) and tangential flow filtration (TFF) allowed for significant depletion of the mostly cell culture supplement-derived EVs. We chose TFF in further experiments for better scalability and time saving purposes because efficient depletion of serum-EVs by UCF required up to 24 h centrifugation at 100,000 x g and was restricted to limited volume^23^. Some but not all tested chemically defined serum-free media contained less than 10^8^ particles/mL (**Fig.S2A**). Culturing PLX cells in fibrinogen-containing α-MEM/HPL resulted in a significant rise of EVs on top of the pre-existing mostly HPL-derived EVs within six days. PLX cell culture in TFF particle-depleted α-MEM*/TFF or in defined media resulted in significantly elevated particle counts after six days indicating effective release of PLX-EVs under these conditions (**Fig.S2B**). Tunable resistive pulse sensing (TRPS) analysis showed a more heterogeneous particle size distribution under serum-free conditions including measurably larger particles (**Fig.S2CD**). For reasons of efficiency, we chose fibrinogen-depleted and particle-depleted α-MEM*/TFF for subsequent experiments.

### Scalable TFF strategy for enriching cell-derived functional EVs

We next devised a scalable EV production process (**Fig.S3**) based on previous results^24^. Fibrinogen-depleted α-MEM* (n x 500 mL) was depleted for preexisting particles (including EVs) before use in large-scale PLX short-term culture. CM was harvested from cell factories after 48 h periods in particle depleted α-MEM*/TFF and subjected to 100x concentration for PLX-EV enrichment (termed EV^TFF1^) and simultaneous soluble factor separation. Isovolumetric washing of EV^TFF1^ removed remaining soluble factors (creating EV^TFF2^).

Cryogenic transmission electron microscopy showed characteristic double membrane-surrounded vesicles (**Fig. 1A**). Super-resolution microscopy of individual EVs confirmed an immunophenotypically heterogeneous EV preparation with co-localization of tetraspanins CD9, CD63 and CD81. Only 11.7% displayed all three tetraspanins simultaneously and 49.8% were single positive for only one tetraspanin (**Fig. 1B**). Western blots confirmed EV identity regarding minimum information for studies of EVs according to MISEV2018 criteria^3^. Comparable CD81 and lower CD9 levels were measured in three representative purified EV preparations from therapeutic lots of PLX cells of three individual placentas. Lineage specificity of tetraspanin expression levels was indicated by lower CD81 and higher CD9 levels detected on endothelial cell-derived control EVs (**Fig. 1C, Fig.S9, Table S1**). Loading corresponding protein amounts showed significantly enhanced tetraspanin and flotillin-1 membrane raft marker signals after sequential TFF. The non-vesicular endoplasmic reticulum lectin, calnexin, and the golgi membrane stacking protein, GRP94, were only found in cell lysates. Apolipoprotein A1 as a marker for plasma-borne high-density lipoproteins was enriched and human serum albumin (HSA) was depleted in TFF2 after enrichment in TFF1 (**Fig. 1C**). Densitometry of EV blots from three independent donors confirmed the results (**Fig. 1D**).

**Figure 1:**
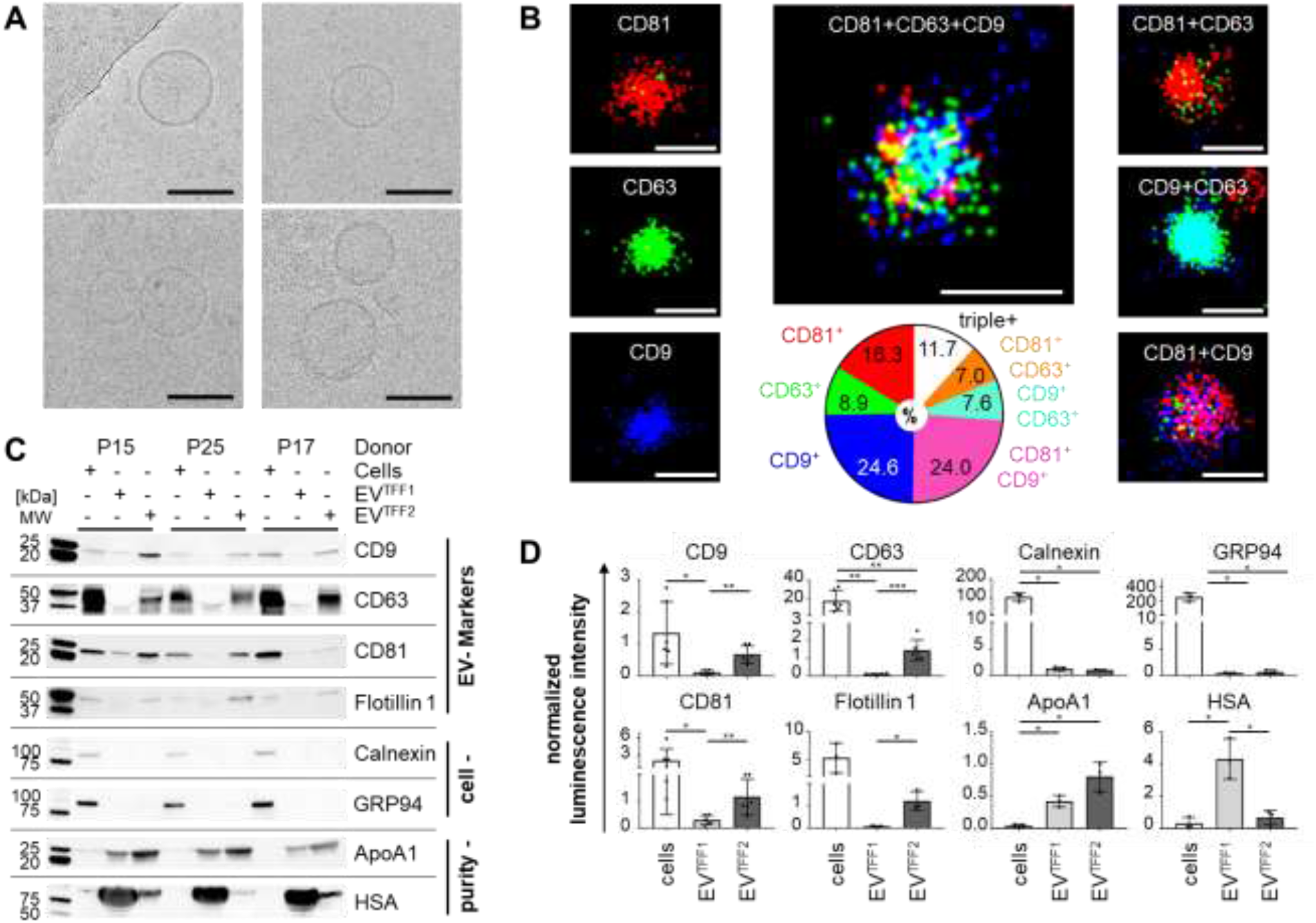
EV identity and purity. (**A**) Cryo-transmission electron microscopy showing representative images of PLX-EVs after TFF (bottom right picture EV^TFF1^, all others EV^TFF2^). (**B**) Super-resolution microscopy topology display of tetraspanins CD81 (red), CD63 (green) and CD9 (blue), via single molecule fluorescence of individual EV^TFF2^ (left) and the overall distribution of double and triple marker positive EVs as indicated in text inserts. (**C**) Western blots comparing placental-expanded (PLX) cells (from lots P15, P25, P27) and corresponding EV preparations after one or two TFF cycles (TFF1, TFF2). Results for tetraspanins CD9, CD63, CD81, EV-specific flotillin, cell markers calnexin and GRP94, as well as culture medium supplement-derived apolipoprotein A1 (ApoA1) and human serum albumin (HSA). (**D**) Densitometry of blots shown in (**C**) normalized to total protein/lane. Significant differences (*p < 0.0332, **p < 0.0021 and ***p < 0.0002) were identified based on two-tailed t-test with 95% confidence level. The complete western blot membranes including also control endothelial cells are shown in **Fig.S9**.

### Trophic proteins regulating angiogenesis and immunity co-enriched with PLX-EVs

Using multiplex bead-based flow cytometry^25^ for EV surface marker profiling confirmed high expression of tetraspanins CD81/CD63 and reduced CD9 expression. We found medium-classified fibronectin receptor CD49e/CD29, high/medium extracellular matrix interaction molecules CD44 and NG2, cytokine receptor CD105 (endoglin) and CD142 (coagulation factor III, tissue factor). The majority of hematopoiesis markers were absent on purified PLX-derived EVs (**Fig.S4A**).

After establishing flow cytometry instrument sensitivity based on silica size marker beads (**Fig.S4Bi**) we defined a gating strategy for nanoparticles sized between ≥ 100 nm up to ≤ 1000 nm. Negative-control phosphate-buffered saline (PBS), with and without calcein and lactadherin reagents, showed only minute unspecific reactivity (**Fig.S4Biii-iv**).

Flow cytometry showed predominantly calcein-converting cell-derived EVs (**Fig.S4Bv**). Calcein and lactadherin signal distribution indicated ≥ 96% EVs derived from intact cells (calcein single-positive) and presumably not apoptotic bodies (lactadherin negative). By back-gating calcein/lactadherin double-positive events (**Fig.S4Bii**, PLX EV size gate plot), a minor proportion of apoptotic bodies was apparent in the 500 – 1000 nm size range as described in the literature^26^ (**Fig.S4CD**).

The EV preparation protocol devised in this study allows for high throughput scalable separation of EVs from soluble factors by sequential TFF processing of CM preparations (**Fig.S5A**). Targeted proteomic pre-analysis using western blot-based arrays comparing unconditioned cell culture media vs. CM soluble factor fractions and TFF1 vs. TFF2-enriched PLX-EVs, respectively, revealed differences between the four fractions (**Fig.S5BC**). The majority of highly abundant proangiogenic factors was present in both, EV^TFF1^ and EV^TFF2^. We did not find specific factors significantly enriched in purified EV^TFF2^ over EV^TFF1^, using this technology (**Fig.S5D**). We next analyzed protein composition of soluble factors and EVs, compared to unconditioned media, to identify proteins uniquely present in different fractions by qualitative label-free proteomics. Pre-analytics confirmed depletion of total protein content in EV TFF fractions by the factor 4.87 – 18.33 accompanied by further 1.34 – 3.30-fold EV enrichment. A total of 401 - 1,168 proteins were detected in different donor-derived fractions. (**Table S2**). Only 708 proteins detected in at least two different donor-derived fractions were selected for further analysis. We identified 258 proteins uniquely present in purified EVs. Of these proteins, 110 were related to either angiogenesis, immune system regulation, cell movement or adhesion. Furthermore, 59 proteins unique to the soluble factors and 18 proteins enriched by TFF were found; 224 proteins were present in all three fractions (**Fig.S5B**). Using differential proteomics with tandem mass tag (TMT) labels, we detected 814 proteins significantly differentially expressed (p < 0.05) with distinct pattern of protein clusters (**Fig. 2A**). Proteins significantly over-represented in EVs after TFF2 related to immune response modulation, angiogenesis, cell movement and EV marker molecules according to selected corresponding GO terms (**Fig. 2B; Table S3**). Ingenuity pathway analysis revealed several canonical protein signaling pathways differentially over-represented in EV preparations compared to soluble factor fractions (**Fig.S6A**). Functional and disease categories related to cell angiogenesis, movement and immune response were classified ‘enriched’ in EV proteomes (**Fig.S6B**).

**Figure 2:**
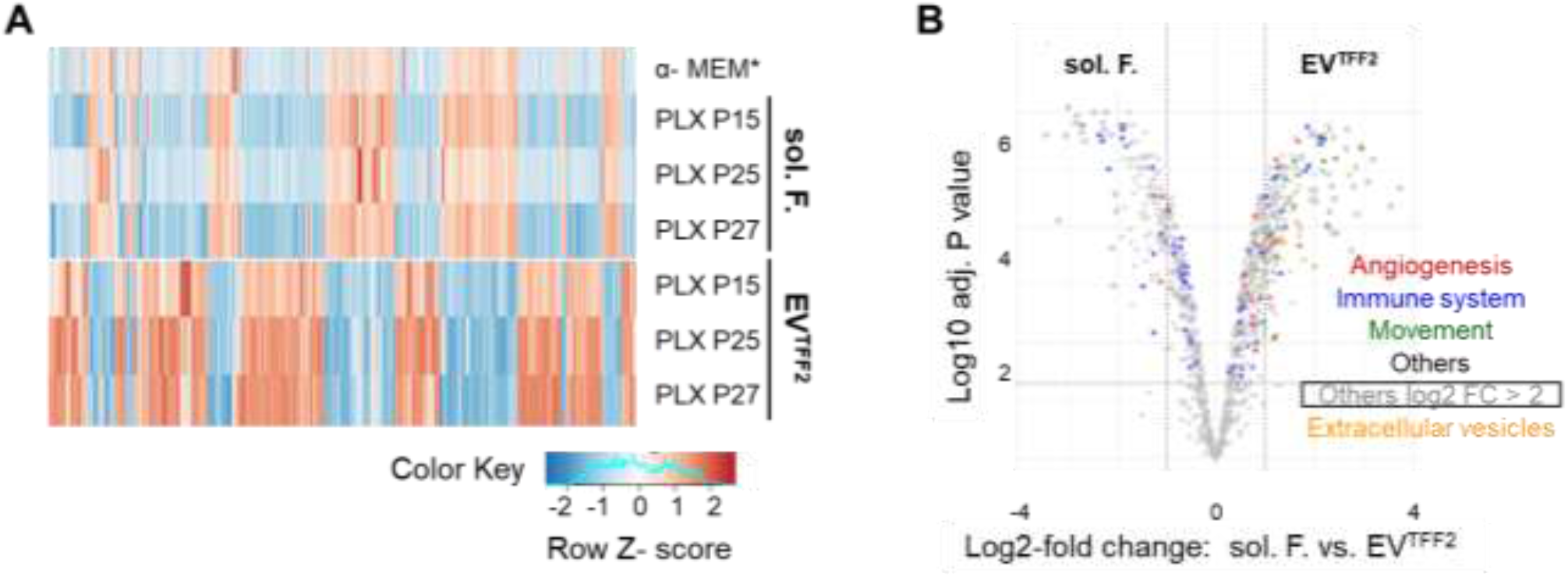
Quantitative proteomics of PLX secretome fractions. (**A**) Heatmap comparing the proteome of soluble factor factors (sol. F.) and EV^TFF2^ with human platelet lysate-supplemented defibrinized unconditioned cell culture medium (α-MEM). (**B**) Volcano plot comparing protein expression signal significance to levels of enrichment in sol. F. vs. EV^TFF2^. Functional categories according to corresponding GO terms as listed in **Table 2** are depicted by color-coded dots as indicated; additional highly over-represented proteins marked as open circles (see also **Figure S6**). Complete tandem mass tag proteomics data were uploaded to PRIDE [accession N°.: PXD014572] including 93 significantly enriched immunomodulatory and angiogenic proteins.

### PLX-EV preparations capture proangiogenic cell-secreted factors and mediate vascularized skin regeneration in vivo

To validate proteomics, we analyzed the different vesicular and vesicle-depleted PLX secretome fractions for their capacity to stimulate endothelial network formation as a surrogate for angiogenesis. EVs and soluble factors efficiently induced network formation in a dose-dependent manner. (**Fig. 3A**). Partial separation of EVs from extra-vesicular proteins by SEC significantly abrogated proangiogenic function of both EVs and soluble factor fractions, indicating EVs gathering trophic factors (**Fig. 3B**). We next hypothesized that the PLX treatment effects could relate to stabilizing therapeutically active proteins in the close vicinity of EVs in a corona-like fashion.

**Figure 3:**
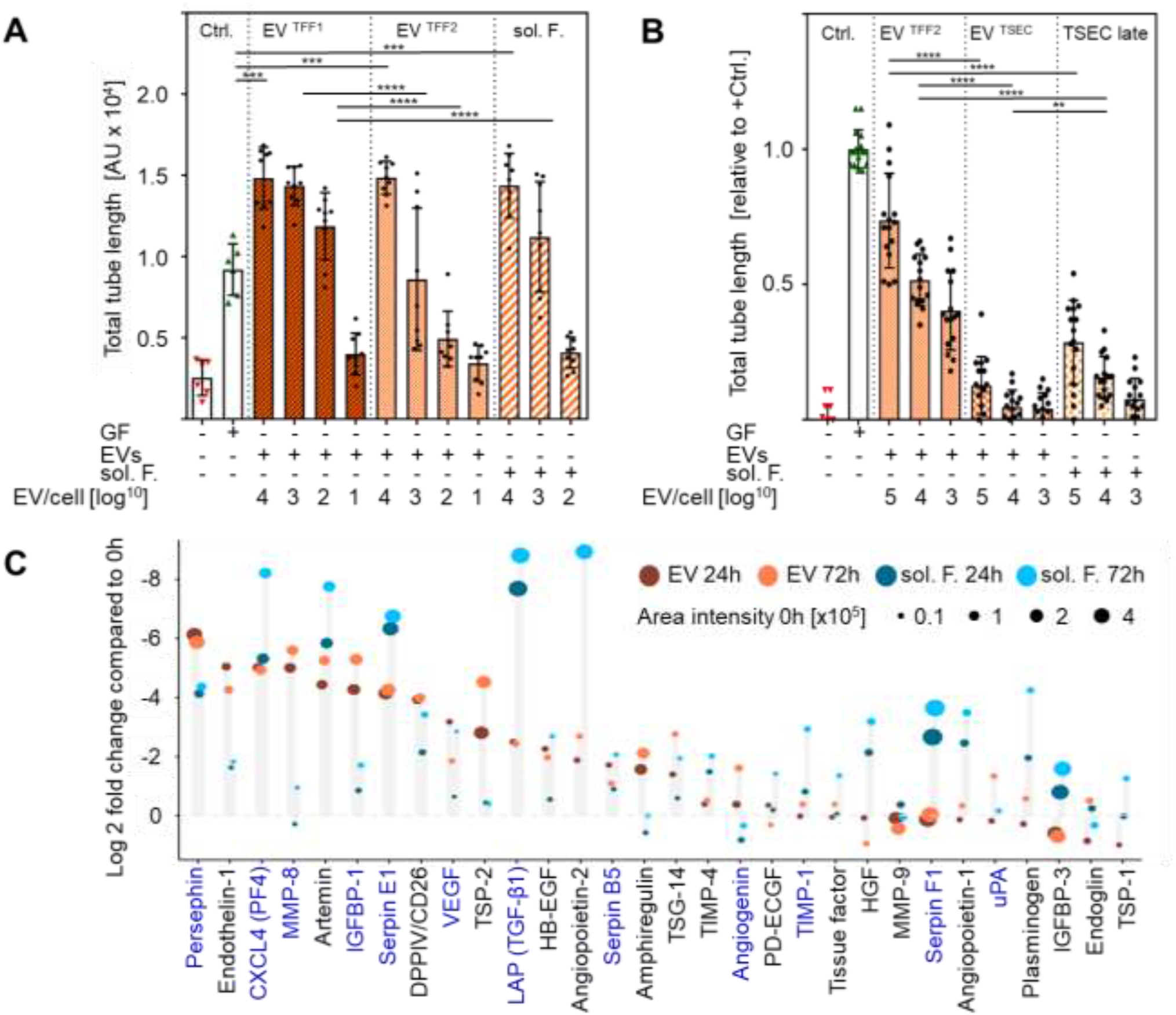
PLX-EVs capture and stabilize proangiogenic factors. (**A**) Angiogenic potential of different PLX secretome fractions as analyzed by endothelial network formation in a matrigel assay. Total length of the endothelial networks in the presence of PLX-EVs at the indicated EV per endothelial cell ratio is shown. Volumes of soluble factors (sol. F.) added to the assay were calculated accordingly corresponding to EV number as described in the methods section. Results pooled from three independent donors (***p < 0.0002, ****p < 0.0001). (**B**) Separation of EV^TFF2^ from their adjacent proteins by size exclusion chromatography (EV^SEC^) revealed significant loss of proangiogenic function. (**C**) Angio-proteome profiling of EVs (reddish dots) vs. sol. F. (bluish dots) after protease treatment, compared to untreated control samples (color code and spot signal area as indicated). T-test followed by Benjamini/Hochberg correction comparing EV^TFF2^ vs. sol. F. preparations after 24 and 72 hours in the presence of 1 mg trypsin/mg protein), respectively. Adjusted p value < 0.05; blue-colored proteins indicating significant protection from degradation in the presence of EVs.

To question in vivo functionality of the trophic factors in EV^TFF2^ preparations, we took advantage of a historical human cell therapy model for deep skin wound regeneration on immunodeficient mice^27^. Using this model, we recently demonstrated that platelet-derived EVs mediate self-assembly of human skin and skin organoids^28^.

Transplantation of human keratinocytes plus fibroblasts in the presence of TFF2-enriched PLX-EVs resulted in rapid human skin regeneration with pronounced early vascularization after two weeks already. EV-deprived PLX-derived soluble factors supported skin regeneration comparable to control transplants. Remarkably, only transplants in the presence of EVs showed patent vessel ingrowth whereas transplants without EVs showed significantly less vessels and pronounced red cell extravasation (**Fig. 4**).

**Figure 4:**
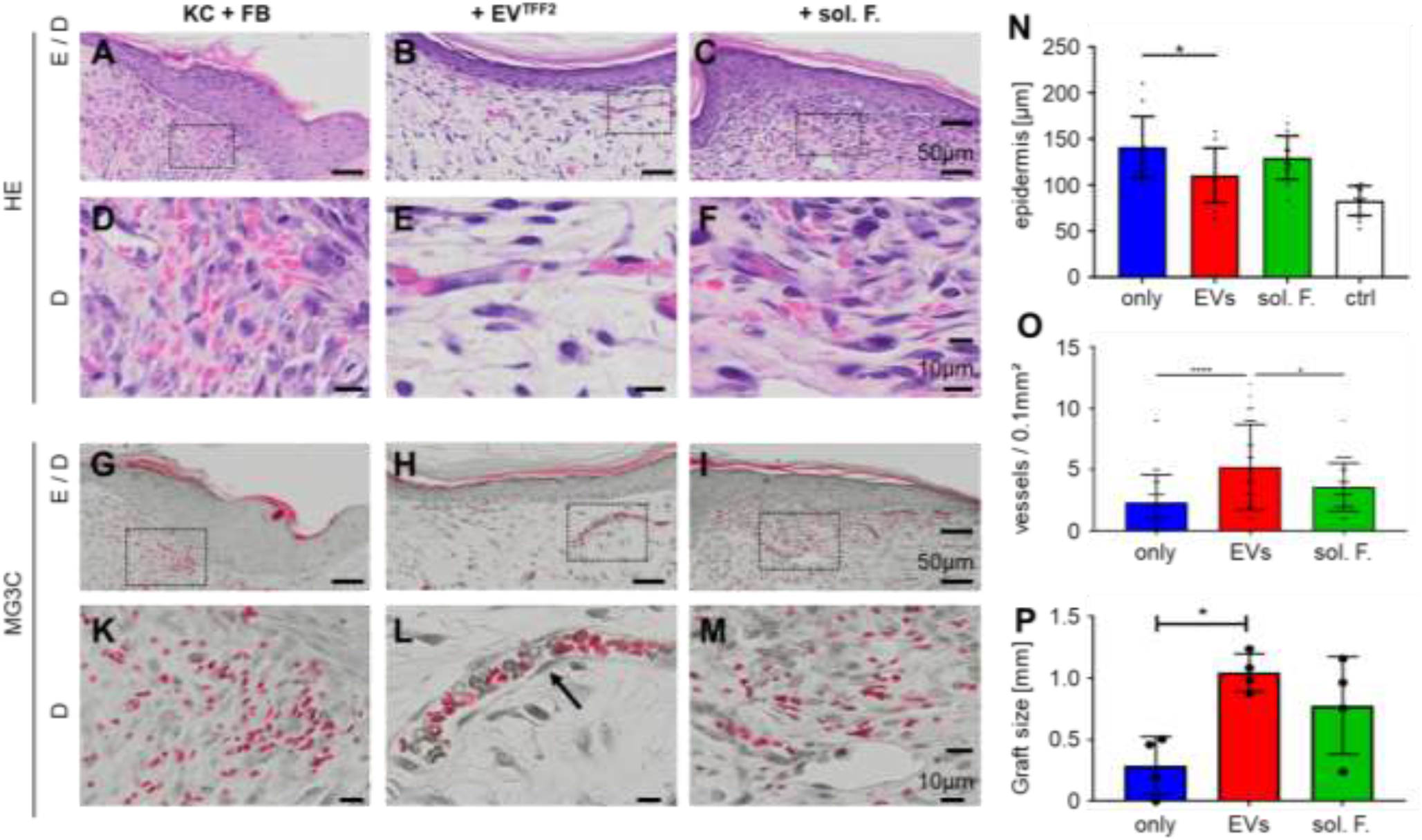
PLX-EVs promote angiogenesis during skin regeneration in vivo. (**A-M**) Histology of keratinocyte + fibroblast (KC + FB) cell suspension grafts in serum, supported by either TFF2-isolated PLX-EVs (+EV^TFF2^) or soluble factors (+sol. F.), day 14 post grafting. Hematoxylin and eosin (HE) and Masson Goldner trichrome (MG3C) staining confirmed the layered cell organization into epidermis and dermis (E/D). Dermal analysis showed vessel enrichment predominantly when supporting the cell grafts with EV^TFF2^. Serum only and sol. F.-driven cell transplants revealed limited murine vessel sprouting and tissue hemorrhage. Vessels were stabilized by pericytes (arrow in L). Dotted boxes in (A-C; G-I) = magnified area in (D-F; K-M). (**N**) Quantification showing significantly increased epidermal thickness in serum only compared to cells grafts supported by EVs^TFF2^. (**O**) Vessel density in grafted dermis and (**P**) graft size was significantly increased in PLX-EV grafts compared to serum only cell grafts^28^. Mean ± SD results (N, O, P). One-Way-ANOVA, multiple comparison of four biological and three technical replicates (*p<0.05, ****p<0.0001).

Engrafted human skin showed appropriate stratification and significantly larger graft size for the EV-treated animals compared to controls (**Fig.S7A-C**). We speculated that extended stability and/or tissue protease resistance of proangiogenic factors in the vicinity to EVs could mediate such differences. To address this issue, we subjected EV^TFF2^ and EV-depleted soluble factors to protease treatment before measuring trophic protein persistence by antibody-based array profiling. EVs protected secreted factors including matrix metalloproteinase-8 (MMP-8), angiogenin, serpins, coagulation factors, angiopoietin and vascular endothelial cell growth factor (VEGF) from degradation (**Fig. 3C**).

### PLX-EV preparations modulate T cell proliferation and target immune cells

To study immunological aspects, we tested the potential of different PLX secretome fractions, compared to parental PLX cells, to inhibit mitogen-driven T cell proliferation. PLX cells and their EVs significantly inhibited T cell proliferation in a dose dependent manner with PLX cells reaching almost 100% inhibition at a 1:1 ratio. PLX-derived soluble factors did not inhibit T cell proliferation in the absence of EVs (**Fig. 5A**). Interestingly, further separating EV^TFF2^ by SEC from their adjacent protein fraction divided the immunomodulatory capacity into EV^TSEC^ and their proteins, both inhibiting T cell proliferation significantly less than EV^TFF2^ (**Fig. 5B**). At a mechanistic level, time-dependent significant uptake of fluorescently labeled EV^TFF2^ was observed in four major immune cell types by multicolor flow cytometry (**Fig. 5C**).

**Figure 5:**
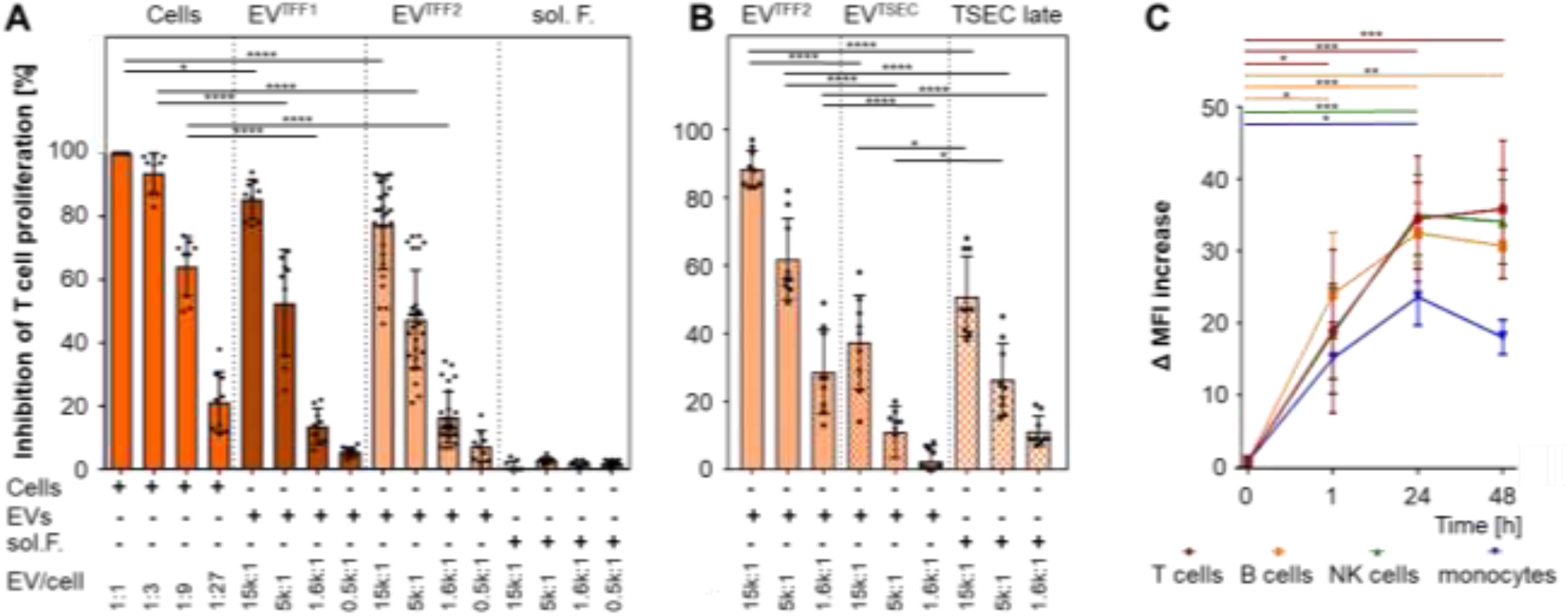
Immunomodulation and cell signaling by EVs: (**A**) Inhibition of phytohemagglutinin (PHA)-induced T cell proliferation by PLX stromal cells, EV^TFF1^ and EV^TFF2^ as compared to soluble factors (sol. F.) added as indicated in limited dilution at defined ratio to mononuclear leukocytes. The percentage of inhibition was calculated relative to the maximum proliferation induced by PHA. Pooled results of three independent donors measured at day four. (**B**). Assay format as used in (A) but testing the inhibition of T cell proliferation by size-exclusion-purified EVs^TSEC^ and their former corona separated by size-exclusion chromatography, compared to parental EV^TFF2^. (**C**) Peripheral blood mononuclear cells were incubated with bodipy-labeled EV^TFF2^ at pre-determined ratio of 1:5,000 for one, 24 and 48 hours as indicated. The bodipy signal was located to CD3^+^ T cells, CD19^+^ B cells, CD56^+^ NK cells and CD14^+^ monocytes by polychromatic flow cytometry. Pooled results from three independent donors performed in triplicate were analyzed (in A-C). Statistical analysis was done in Graph-Pad Prizm 7.03 using one-way Anova analysis with Sidak correction for multiple samples (** p < 0.002; *** p < 0.0002; **** p < 0.0001).

Confocal microscopy of sort-purified cells confirmed cytoplasmatic localization of EV^TFF2^ after 24 h (**Fig.S8A**). Sort-purified T cells and monocytes were selected to determine intracellular signaling of EVs in target cells. A differential signaling signature was observed with over-representation of phosphorylated cyclin-dependent kinase inhibitor p27 and glycogen synthase kinase-3, regulating cell cycle and signaling, respectively (**Fig.S8B**).

### Evidence for the EV corona

To address the question whether a protein corona determines EV function, as observed previously for various synthetic and inorganic nanocarriers, we compared EV^TFF2^ (i.e. corona-bearing) vs. ultracentrifugation-purified EVs (i.e. corona-depleted) for their proangiogenic potential. Ultracentrifugation resulted in significant loss of EV-mediated vascular network formation. We selected VEGF, insulin-like growth factor-1 (IGF) and epithelial growth factor (EGF) dissolved in albumin solution (VIE/A) for re-establishing an artificial corona around ultracentrifugation-purified EVs. The VIE/A-cloaked EVs significantly re-established their proangiogenic potential in a corona protein-dose-dependent manner. These corona-bearing EVs were significantly more efficient than respective doses of VIE/A in the absence of EVs over a large log-range of concentrations. Adding human serum albumin to EVs without VIE led to minor but significant improvement of EV function (**Fig. 6AB**). Significant 10-fold relative depletion of VEGF, as reference angiogenesis factor, was confirmed by enzyme-linked immunoassay after ultracentrifugation of EV^TFF2^ (**Fig. 6C**).

**Figure 6:**
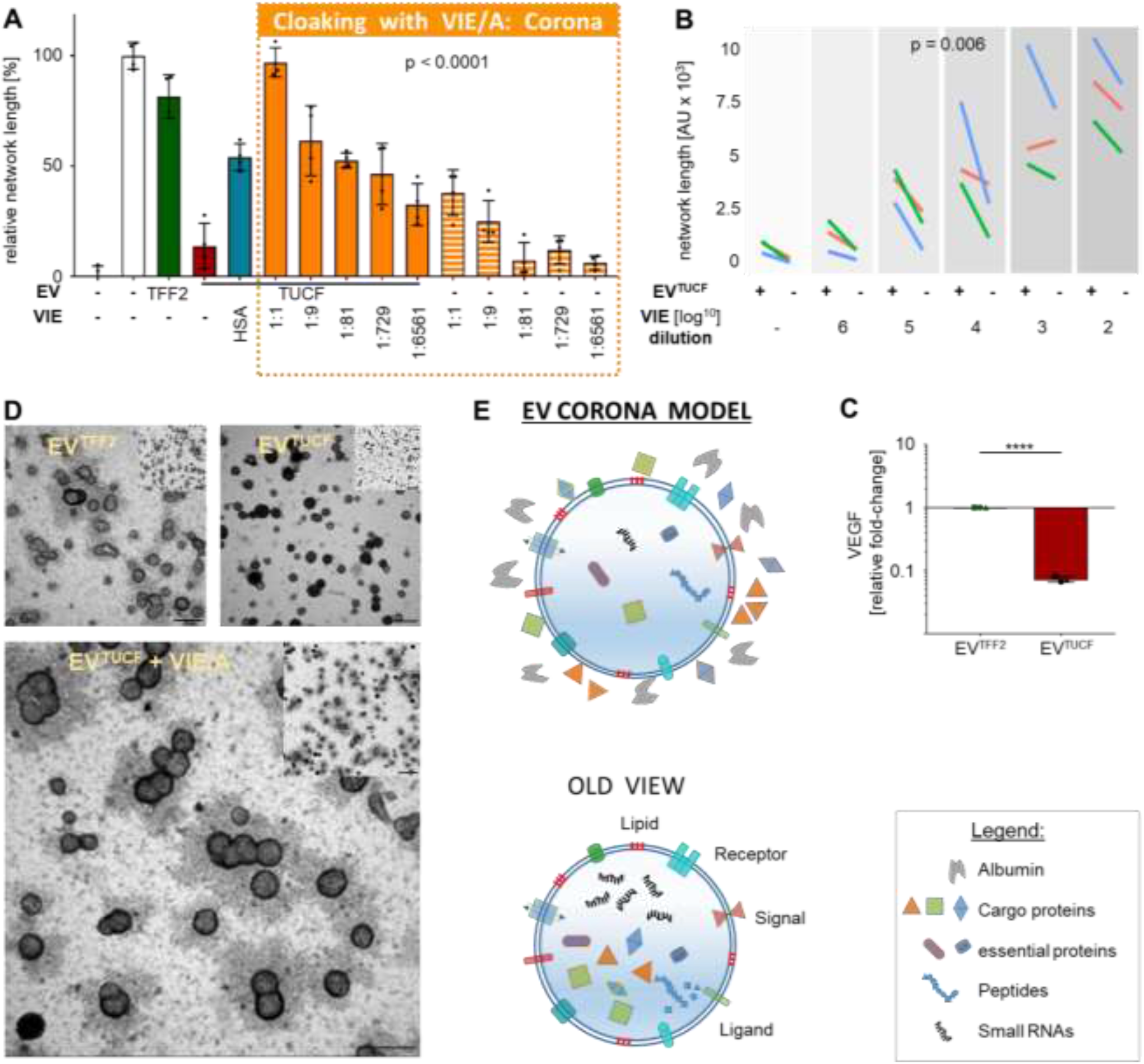
EV protein corona location: (**A**) Endothelial cell network formation comparing EVs^TFF2^ with ultracentrifuged corona-depleted EVs (EV^TUCF^) and free growth factors and growth factor corona-cloaked EVs. Representative experiment, quadruplicates. (**B**) Dose dependent endothelial network formation in the presence (+) or absence (-) of EV^TUCF^ in an 8% albumin solution containing VEGF, IGF and EGF (VIE/A) (n = 3; individual additional experiments color-coded in red, green, blue, and covering a broader dose range than in (A),). Statistics were calculated using a linear mixed model with treatment as fixed effect and dilution as random effect (A, B). (**C**) Fold-change of VEGF concentration comparing EVs^TFF2^ to EVs^TUCF^ (mean ± SD, n = 3, T-test, p < 0.05). (**D**) Negative contrast transmission electron microscopy images of PLX EVs purified with tangential flow filtration (TFF2), depleted protein corona after TFF combined with ultracentrifugation and with protein corona reestablished by defined factors VEGF+ IGF+ EGF in albumin solution (VIE/A). Scale bar 250 nm, small insert showing overview image with scale bar 500 nm. (**E**) New EV corona model and current/old view on EV structure (modified from^30^ and www.exosome-rna.com/evpedia).

To visualize the postulated protein corona around PLX-EVs we took advantage of conventional negative-contrast transmission electron microscopy. EVs^TFF2^ showed a halo around vesicles, suggestive of a corona, that was absent after ultracentrifugation. The halo appeared again after cloaking ultracentrifugation-purified EVs with VIE/A indicating corona formation (**Fig. 6D**). These pictures were highly reminiscent of previous results showing the prototypic polystyrene nanocarrier corona^29^. Based on these results we suggest a new model that considers the nanoparticle corona to be also operative on cell-derived EVs (**Fig. 6E**).

## DISCUSSION

In this study, we devised an efficient scalable isolation and purification strategy for mass-production of EVs enabling high-content proteomic and functional analysis. Sequential TFF concentration enriched EV-adjacent proteins in a fashion reminiscent of the protein corona that conventional nanoparticles acquire in biologic fluids^31^. Based on previously established large-scale cell propagation protocols^32^, up to one trillion EV^TFF2^ were obtained from 1.5 liter CM within less than a working day (e.g. 5 - 10 mL final TFF2 concentrate including 1 - 2 x 10^11^ EVs/mL). TFF-based particle depletion of starting media or the use of defined particle-poor media permits directed cell-derived EV analysis devoid of serum- or culture supplement-derived EV or particle contamination. EV identity and purity achieved with this strategy was confirmed by western blot and flow cytometry according to MISEV 2018 criteria^3, 33^.

Taking advantage of the straightforward separation of soluble factors from EVs out of CM by stepwise TFF (**Fig. S3**), we were able to perform in-depth proteomic analysis. We demonstrate that proteomes in soluble factor as compared to EV fractions differ significantly, compatible with an enrichment of certain proteins in the vicinity of EVs. Despite these differences, both TFF-purified PLX-EV fractions as well as the soluble factors induced supra-maximum vascular network formation in a dose dependent manner *in vitro*. This could be explained by the fact that several proteins involved in vascular remodeling, e.g. serpins, platelet factor-4 (PF-4), thrombospondin-1 (TSP-1) and angiogenin, were present in the soluble factor fraction as well as in EV preparations. Provided, that a comparable ‘soluble factors plus EVs’ secretome pattern also occurs in patients, our results would indicate that PLX cell-derived EVs may be considered to contribute to the therapeutic effect. We may speculate that EVs can act over longer periods of time beyond rejection of the allogenic cell therapy and degradation of secreted factors. A precise assignment of the various effects rendering different aspects of vascular and tissue remodeling and wound healing induced by PLX cells or their soluble and vesicular secretome fractions requires additional research that is currently underway in mechanistic side studies accompanying the ongoing clinical trial (http://www.pace-h2020.eu/).

The canonical view on cell-derived EVs displays a cargo-carrying core enveloped by a receptor-bearing double membrane (**Fig. 6E**). This basic structure was sufficient to explain many EV functions during cell-to-cell communication and built the basis for designing specialized EV products as next-generation drug delivery carriers^19^. Our observation that cell-derived EVs can acquire a functional corona, together with our demonstration that creating an artificial corona by cloaking ultracentrifugation-purified EVs with VIE/A can re-establish proangiogenic function, adds another level of complexity to our understanding of EV biology. The multiple facets of nanoparticle corona functions are well established. Our study highlights that also cell-derived EVs can be subject to corona formation. Additional corona protein compositions may be derived from proteomics. Also, the precise molecular mechanisms leading to release of the functional EV corona components at the target endothelial cell needs to be disclosed. We can just speculate whether corona-bearing EVs can also form a nanoscale synapse, as proposed for other nanostructures^34^, when executing their function. It was also interesting to note, that the PLX-EVs could protect certain corona proteins from protease-mediated degradation.

We identified immunity in addition to angiogenesis as predominantly affected by pathways that require action of the proteins over-represented particularly in purified PLX-EVs. PLX, like other stromal cells, have immunomodulatory capacity^15^. In contrast to angiogenesis, dose-dependent inhibition of T cell proliferation was observed only with PLX cells and their EVs. Understanding the relative contribution of cells vs. soluble factors vs. EVs to therapeutic efficacy will be a major challenge on the way towards rational design of efficient cell-derived and EV-based therapies. This study has certain limitations. The precise position of EV-associated proteins still needs to be determined. Questions regarding soft vs. hard corona composition and function need to be addressed. The continuous enrichment of corona-bearing EVs with soluble protein reduction throughout the purification process argue in favor of TFF-based protocols as a baseline for these studies. Reaching a higher level of purity without affecting the corona is expected to result in more precise information during future high content omics screening.

The idea of an EV corona has implications for both nanoengineering of EV-inspired synthetic nanocarriers and EV-based therapy development.

### MATERIALS & METHODS

All materials and methods used in this study are summarized in detail in the online section.

## ONLINE METHODS SECTION

### Ethics statement

Animal trial permission was given by local authorities according to Austrian legislation (§ 26 TVG 2012; animal trial number: BMBWF-66.019/0032-V/3b/2018). Human full-thickness skin was obtained as biological waste material after informed consent as approved by the ethical committee of the region of Salzburg (vote number: 415-E/1990/8-216).

### Cell culture media and reagents

Serum- or plasma-containing media tested in this study included α-MEM, Eagle’s MEM (EMEM, both Sigma-Aldrich, USA) and endothelial growth medium (EGM2, Lonza, USA). Media were supplemented with 10% pooled HPL or FBS as indicated, 5 mM N(2)-L-alanyl-L-glutamin (Dipeptiven, Fresenius Kabi, Austria) in the absence or presence of 2 U/mL preservative-free heparin (Biochrom, Germany) without antibiotics or with 100 U/mL penicillin and 0.1 mg/mL streptomycin (both Sigma-Aldrich, USA) as indicated in the results section. For endothelial cell culture, EGM-2 was supplemented with hydrocortisone, fibroblast growth factor-2 (FGF-2), VEGF, IGF, epidermal growth factor (EGF) and ascorbic acid from the supplied bullet kit (all Lonza) and FBS was replaced by an equivalent volume of pHPL^35^. For particle depletion, media were prepared by clotting supplemented α-MEM as described without heparin to avoid possible inhibition of EV function^36^. The collapsed fibrin clot was removed by centrifugation for 10 min, 3,000 x g, at room temperature. The resulting pre-cleared medium was filtered through a 0.22 µm stericup filter (Merck Millipore, USA) and finally particle depleted using a 1,600 cm² 500 kDa cut off hollow fiber modified polyethersulfone (mPES) membrane filter column operated on a KR2i TFF System (Repligen, USA). This medium was termed α-MEM*/TFF. For comparison to TFF, we used ultracentrifugation that was performed at 100,000 x g for 3 h in a Sorval WX80 ultracentrifuge with a T-865 rotor (both Thermo Scientific, USA) as indicated.

Chemically defined and serum-free media included CNT-Prime (CELLnTEC, CH), X-Vivo-10, -15 and -20 (Lonza, USA) and MSC NutriStem XF medium (including a proprietary supplement mix; Biological Industries, Israel). Media were 0.22 µm sterile filtered prior to use.

### PLX cell culture

Cryopreserved clinical grade PLX cell aliquots were obtained from Pluristem Ltd. (Israel) as part of a European funded research project (http://www.pace-h2020.eu/). For this study, we used PLX cells derived from three individual donors (P150216R01, P250416R05 and P270114R27, termed P15, P25 and P27, respectively). For EV production, aliquots of 2.5 million PLX cells were propagated for one to two passages to avoid excess proliferation at moderate seeding density of 1,000 cells/cm² in 2,528 cm² cell factories (CF4, Thermo Fisher Scientific, USA) as established previously until approximately 70% confluence in α-MEM/10% HPL^37, 38^. For EV harvest, PLX cells were washed twice with 37°C pre-warmed Dulbecco’s phosphate buffered saline (PBS, Sigma Aldrich, USA) and cultured in particle depleted α-MEM*/TFF for one to three additional 48 h periods with complete CM change.

### EV quantification by TRPS

We used TRPS to quantify the particle content in various samples including fresh cell culture media, conditioned media, crude and TFF2 EV preparations. Samples were diluted at least 1:1 in Dulbecco’s PBS containing 0.05% Tween 20 that was also used as measurement electrolyte. Measurements were performed using a qNano Gold (Izon, New Zeeland) equipped with an NP150 nanopore (analysis size range 70 - 420 nm).

### EV enrichment by TFF and purification by size exclusion chromatography (SEC)

To isolate and purify EVs from particle-depleted CM, the conditioned medium was harvested after 48 h intervals. Possibly remaining cells were depleted by centrifugation at 300 x g for 5 min followed by a 3,000 x g centrifugation step for 10 min to deplete cell debris. This pre-cleared conditioned medium was first concentrated 100-fold using a 300 kDa cut off hollow fiber mPES membrane filter column operated on a KR2i TFF System (Repligen, USA). The particle-free soluble factor fraction was collected as permeate at this step while the TFF1 preparation was kept as ‘retentate’ inside the system. By washing this TFF1 fraction with twice the starting volume NaCl 0.9% buffered with 10 mM HEPES an additional depletion of proteins and other non-vesicular non-particulate content was obtained, resulting in more purified but still protein-containing EV^TFF2^ fraction. SEC was performed using qEV 70 columns (Izon, New Zealand) according to manufacturer’s instructions to further purify EVs and remove extra-vesicular putative corona protein (**Fig.S3**). EVs were pooled from early fractions (7 - 9) whereas late fractions (17 - 19) were pooled as comparative corona-derived protein samples.

### EV corona removal with ultracentrifugation and re-establishment of a defined EV corona

To completely remove the protein corona from EV^TFF2^ preparations they were diluted 1:10 with NaCl 0.9% buffered with 10 mM HEPES and pelleted via ultracentrifugation at 4°C with 110,000 x g for 90 min in a Sorval WX80 ultracentrifuge with a TH-641 rotor at 25,400 rpm all (Thermo Fisher, USA). The resulting pellet was re-suspended in the initial sample volume with NaCl 0.9% buffered with 10 mM HEPES. To re-establish a protein corona on the naked EV^TUCF^ they were incubated for 1h at 37°C in EGM-2 (Lonza) 4 % of human serum albumin (Fresenius Kabi, Austria) and 1:100 dilution of VEGF, IGF and EGF (Lonza) or a 1:10 dilution as indicated in the figure legends.

### EV identity analysis by western blot and flow cytometry

To analyze EV identity and purity aspects, we performed western blotting using TGX stain free gradient 4 - 20% SDS-PAGE gels run in a mini-Protean system. Samples were loaded with Laemmli buffer containing 50 µM dithiothreitol (DTT), as reducing agent, except for tetraspanins CD9, 63 and 81. After transfer using the mini Trans-Blot tank system (all Bio-Rad, USA), nitrocellulose membranes were probed with primary antibodies diluted in Tris buffered saline with Tween 20 detergent (TBST) including 2% bovine serum albumin (BSA) in the dilutions indicated in **Table S4**.

Detection was performed using horseradish peroxidase (HRP)-labeled secondary antibodies (rabbit anti-mouse IgG, A27025, Thermo Fisher, USA; mouse anti-goat 205-035-108, Jackson Laboratories, USA; or polymer goat-anti-rabbit, K4002, DAKO EnVision, Agilent, USA) depending on the host species of the primary antibody and clarity enhanced chemiluminescence (ECL) substrate. Bands were visualized and quantified using a chemidoc system and image lab software (all Bio-Rad). Densitometry of specific bands was quantified after background correction in relation to the total protein content detected with the stain free technology before transfer.

In order to obtain a broader overview of markers present on the surface of PLX-derived EVs we applied a bead-based screening assay measured with flow cytometry as described^39^. To standardize the EV input for the assay, we loaded 1 x 10^9^ EVs^TFF1/TFF2^ on the MACS Plex capture bead mix (Miltenyi, Germany)^25^ stained with a mix of CD9-, CD63- and CD81-APC detection reagent mix according to manufacturer’s protocol. Analysis was performed with an LSR Fortessa instrument (BD, USA) equipped with 355 nm, 405 nm, 488 nm, 561 nm and 640 nm lasers. Raw measurement data were corrected for unspecific binding of the detection antibody mix to beads and expressed as relative fold-change of mean fluorescence intensity (MFI) compared to samples stained with isotype control.

For single EV analysis by flow cytometry EV ^TFF1^ and EV^TFF2^ preparations were stained with lactadherin-Alexa fluor 647 (CellSystems, Germany), and calcein AM (Sigma-Aldrich, USA) that were aggregate-depleted by centrifugation at 17,000 x g for 10 min directly prior to use. Staining with lactadherin was performed for 30 min at 4°C and subsequently, after 1:10 dilution of the samples, with calcein for 5 min at room temperature. Samples were further diluted 1:5 in PBS and analyzed on a Cytoflex flow cytometer (Beckman Coulter, USA). For creating a size-based gate < 1000 nm for EV detection, we used 100, 200, 500 and 1000 nm green fluorescent-labeled silica beads (Kisker Biotech, Germany) as illustrated in **Figure S4B**. Based on the separation of fluorescent EVs from particle background we only recorded fluorescence-positive events using double-fluorescence triggering in the FITC and APC channels. For quantifying calcein^+^ and lactadherin^+^ EV, gating was performed using staining reagents appropriately diluted in PBS as a negative control. Percentage of calcein, lactadherin and double-positive populations was calculated as a part of the sum of all acquired events and were displayed as a pie chart (**Fig.S4C**).

### Electron microscopy

For conventional negative contrast transmission electron microscopy (TEM), 10 µL EV samples were applied on formvar-coated 100 mesh copper grids (Agar scientific, UK), fixed with 2.5% glutaraldehyde and stained with uranyl acetate replacement solution 1:10 (Electron Microscopic Sciences, UK) in bi-distilled water. Dried samples were imaged using an 80 kV LEO EM 910 transmission electron microscope (Zeiss, Germany) equipped with a Tröndle 227 Sharp Eye digital camera system. For cryo-TEM, EV samples were diluted 1:10 in 0.9% NaCl solution and 4 µl were applied to Quantifoil (Großlöbichau, Germany) Cu 400 mesh R1.2/1.3 holy carbon grids (Leica Microsystems, Germany). Grids were glow discharged for 1 min at -25 mA with a Bal-Tec (Balzers, Liechtenstein) SCD005 glow discharger and loaded into a Leica GP grid plunger with the climate chamber set at 4°C and 70% relative humidity. EV samples were diluted 1:10 in 0.9% NaCl solution and 4 µl were applied to the carbon side of the grid. After front-side blotting for 2 - 8 seconds (using the instrument’s sensor function, no pre- or post-blotting incubation) with Whatman filter paper #1 (Little Chalfont, Great Britain) grids were plunge frozen into liquid ethane at approximately -180° C for instant vitrification. Cryo-samples were transferred to a Glacios cryo-transmission microscope (Thermo Scientific, USA) equipped with an X-FEG and a Falcon III direct electron detector (4,096 x 4,096 pixels). The microscope was operated in a low-dose mode using the SerialEM software^40^. Images were recorded digitally in linear mode of the Falcon III camera at magnifications of 5,300 (pixel size: 27.5 Å, defocus: -50 µm, dose: 0.2 e/Å2), 36,000 (pixel size: 4.1 Å, defocus: -6 µm, dose: 14 e/Å2) and 150,000 (pixel size: 0.98 Å, defocus: -3 µm, dose: 60 e/Å2).

### Super-resolution microscopy

For super-resolution analysis of EVs, samples were immobilized on microfluidic glass slides (EV profiler Kit, Oxford Nanoimaging, UK) and stained with CD9-ATTO488, CD63-Cy3 and CD81-Alexaflour 647 according to manufacturer’s instructions. Images were acquired via direct stochastic optical reconstruction microscopy (dSTORM; Nanoimager S, Orford Nanoimaging, UK) using 30%, 40% and 50% power on the 488 nm, 561 nm and 640 nm laser, respectively. 2,500 images per channel were recorded for localization mapping. Co-localizations of the different tetraspanins were analyzed using the CODI platform (https://alto.codi.bio/). Localization clusters showing more than 10 individual localizations were considered as EVs. EVs were considered positive for a marker when more than 5 individual localizations were detected in the same channel in a radius of 90 nm around the center of a cluster. For statistical analysis, three fields of view were acquired for three independent donors.

### Proteomics and bioinformatics

Reagents included acetonitrile (≥ 99.9%) and methanol (≥ 99.9%; both VWR, Austria), DTT (≥ 99.5%), formic acid (98 - 100%), iodoacetamide (≥ 99.0%), sodium dodecyl sulfate (SDS; ≥ 99.5%), triethyl ammonium bicarbonate (TEAB, 1 mol/L) and trifluoroacetic acid (≥ 99.0%; all Sigma-Aldrich, Austria), ortho-phosphoric acid (85%; Merck, USA), and sequencing grade modified trypsin (Promega, USA). Deionized water was purified with a MilliQ^®^ Integral 3 instrument (Millipore, USA).

To determine protein content, samples were adjusted to 5% SDS and 50 mmol/L TEAB (pH 7.55) and incubated at 95°C for 10 min to lyse the EVs. Samples were analyzed by a Pierce™ bicinchoninic acid protein assay kit (Thermo Fisher Scientific, Austria) according to the manufactureŕs instructions. S-Trap™ mini columns (Protifi, Huntington, NY, USA) were utilized for sample preparation and 100 µg of protein were prepared according to the manufactureŕs instructions with minor adjustments: Lysis of EVs as well as denaturation and reduction of proteins were performed in 5% SDS and 50 mmol/L TEAB supplemented with 40 mM DTT at 95°C for 10 min. Cysteines were alkylated by the addition of IAA to a final concentration of 80 mM and incubated in the dark for 30 min. Proteins were digested within the S-Trap matrix with trypsin at an enzyme to substrate ratio of 1:10 w/w at 37°C for 18 h. Peptides were eluted and subsequently dried using a vacuum centrifuge. These samples were re-suspended in H_2_O + 0.1% FA to a concentration of 3.33 mg/mL for analysis in a label-free manner. Peptides of each sample (20 µg) were labeled by a TMT 10-plex™ kit (Thermo Fisher Scientific, Austria). Labeled samples were pooled and desalted using 100 µL Pierce™ C18 tips (Thermo Fisher Scientific, Austria) and dried again using a vacuum centrifuge. These samples were re-suspended in H_2_O + 0.1% FA to a concentration of 5 mg/mL.

High-performance liquid chromatography (HPLC) separation was carried out on a nanoHPLC instrument (UltiMate™ U3000 RSLCnano, Thermo Scientific, Germany) at a flow rate of 300 nL/min and a column oven temperature of 50°C. Separation of unlabeled samples was performed on an Acclaim™ PepMap™ 100 C18 column (500 mm x 75 µm i.d., 3 μm particle size, Thermo Fisher Scientific, Austria). Samples (3.3 mg/mL; 0.15 µL) were injected using a microliter pick-up mode (loop volume 1 µL). A multi-step linear gradient of mobile phase solutions A (H_2_O + 0.1% formic acid) and B (acetonitrile + 0.1% formic acid) was applied as follows: 1% - 22% B for 200 min, 22% - 30% B for 40 min, 30% - 55% for 30 min, 90% B for 20 min and 1% B for 40 min. Each sample was measured once.

Separation of tandem mass-tag (TMT)-labeled samples was performed on a 2,000 mm µPAC™ C18 column (PharmaFluidics, Ghent, Belgium). The sample [5.0 mg/mL; 1µL] was injected using a microliter pick-up mode (5 µL loop volume). A multi-step linear gradient of mobile phase solutions A and B was applied as follows: 1% - 22% B for 500 min, 22% - 40% B for 100 min, 90% B for 30 min and 1% B for 100 min. Five technical replicates were measured.

All mass spectrometry measurements were conducted in positive ion mode on a hybrid mass spectrometer (QExactive™ Plus benchtop quadrupole-Orbitrap® mass spectrometer) equipped with a Nanospray Flex™ ion source (both Thermo Scientific, Germany) and a SilicaTip™ emitter with 360 µm outer diameter, 20 µm inner diameter, and a 10 µm inner tip diameter (New Objective, Woburn, MA, USA). Mass spectrometric data were acquired with the following instrument settings: spray voltage of 1.5 kV, capillary temperature of 320°C, S-lens, radio frequency level 55.0, MS1 AGC target 3 x 10^6^, m/z range 400 – 2,000, maximum injection time of 100 ms, resolution of 70,000 at 200 m/z. Data-dependent tandem mass spectrometry (ddMS2) was carried out in the higher-energy collisional dissociation (HCD) cell at a normalized collision energy (NCE) setting of 28.0 and a resolution setting of 17,500 at m/z 200 for unlabeled samples and at a resolution of 35,000 at m/z 200 for TMT-labeled samples. For MS2, the top 15 signals were chosen for fragmentation with a 2.0 m/z isolation window, an automatic gain control and maximum injection time of 100 ms. The dynamic exclusion was set to 30 seconds. The instrument was calibrated using Pierce™ LTQ Velos ESI Positive Ion Calibration Solution (Life Technologies, Vienna, Austria).

All data were evaluated using MaxQuant software (version 1.6.1.0) using default settings. A protein list was obtained from the Uniprot database including both Swiss-Prot as well as TrEMBL entries for homo sapiens (access: 10.03.2019) and was provided for MaxQuant searches^41, 42^. TMT-labeled data were further processed using Perseus software package (version 1.6.1). Analysis of the TMT-labeled samples was conducted using Ingenuity^®^ Pathway Analysis (IPA; version 47547484; Qiagen Bioinformatics, Redwood City. CA, USA). R software (www.R-project.org) was used all further proteomics analysis. For the TMT proteomics data, contaminants were removed and values were log2 transformed and normalized by subtraction of the median of each channel. In order to see how samples cluster together, a principle component analyses (PCA) and hierarchical clustering analysis using Euclidean distance were conducted on the whole normalized dataset. Then, we conducted differential expression analysis using limma package and p-values were corrected using Benjamini and Hochberg multiple testing correction. Proteins were considered significantly differentially expressed if the corrected Benjamini and Hochberg p-value was < 0.05 and absolute log2 fold change > 0.6. Data from the label-free proteomics analysis were used in order to detect proteins present in specific fractions. Samples were considered detected in a specific fraction if they were present in all three replicates. In order to estimate the biological process gene ontology (GO) terms enriched in the purified EV fraction, compared to the soluble factors, we combined the proteins that were significantly enriched in the EV fraction compared to the soluble factors (in the TMT proteomics analysis) and found only in the EV fraction using the label-free proteomics analysis. Enrichment analysis was conducted using ClusterProfiler R package and GO were considered significantly enriched if the adjusted p-value was < 0.01 and the gene count was > 5. Only the most significantly enriched proteins were shown (fold enrichment > 4).

### Angiogenesis and immunomodulation

To assay angiogenic potential of different PLX secretome fractions we used a vascular–like network formation assay on matrigel as previously described ^32^. Umbilical cord blood (UCB)-derived ECFCs were seeded on top of matrigel (angiogenesis assay kit, Merck Millipore, USA; **Fig. 3A**) or a reduced growth factor basement membrane matrix (Geltrex, Thermo Fisher, USA; **Fig. 3B**) in an angiogenesis 96-well µ-plate (Ibidi, Germany) at a density of 31,500 cells/cm^2^. Cells were treated with crude or purified EV preparations in an EV to ECFC ratio of 10,000:1, 1,000:1, 100:1 and 10:1, or with the volume equivalent to EV-free PLX stromal cell-derived soluble factors. Completely supplemented EGM-2 (Lonza) served as a positive control and nonsupplemented EBM-2 with 2 or 4 % of human serum albumin (Fresenius Kabi, Austria) as a negative control. Images were taken every hour for 12 hours on an Eclipse Ti inverted microscope (Nikon) equipped with a custom-build live cell incubation system (Oko Lab, Italy and Nikon, Austria) using a 4x objective. Images were processed with the NIS Elements Advanced Research package analysis software (Nikon). Total matrigel areas were cut out of raw images, homogenized (strength 16), and subjected to intensity equalization. Afterwards, pictures were sharpened slightly and denoised (advanced denoising 5.0). Finally, lookup tables were adjusted to 3,000-13,000 and images were exported as TIFF files. Exported images were cut at the diameter of 1,300 pixels to remove edges of the plate and the contrast was enhanced. Processed pictures were analyzed to automatically detect the network structures with Image J using the Angiogenesis Analyzer plugin and total length of tube-like structures was determined (https://imagej.nih.gov/ij/macros/toolsets/Angiogenesis%20Analyzer.txt).

The effect of PLX-derived EVs on the immune response was analyzed using a T cell proliferation assay as published earlier ^24, 43^. In brief, peripheral blood mononuclear cells (PBMCs) were isolated and pooled from 10 individual donors before labelling with carboxyfluorescein succinimidylester (CFSE; Sigma-Aldrich, USA) and cryopreservation in appropriate aliquots for later use. These pre-labeled PBMCs (300,000 per flat-bottomed 96-well plate) were than stimulated with 5 µg/mL phytohemagglutinin (PHA, Sigma-Aldrich, USA) to induce mitogenesis (at day four). Assays were once loaded with EV doses at three-fold serial dilution in a ratio of 15,000:1, 5,000:1, 1,666, or 555:1. The percentage of proliferating T cells was measured by flow cytometry as the fraction of viable CD3 positive cells with reduced CFSE staining compared to non PHA-stimulated cells. Inhibition of T cell proliferation was expressed as percentage relative to maximum proliferation without EV addition.

### Proteolytic stability and VEGF immunoassay

To investigate the proteolytic stability of proangiogenic and immunomodulatory factors in EV preparations compared to soluble factors, samples were mixed 1:1 with cell culture grade 0.25% trypsin solution (Gibco, Thermo Fisher, USA) and incubated as indicated. To block remaining trypsin, activity reaction was stopped by adding Halt protease inhibitor cocktail or 1mM phenylmethylsulfonyl fluoride serine protease inhibitor (Thermo Fisher, USA). VEGF concentration in EV^TFF2^ vs. EV^TUCF^ was measured in duplicates for each sample (n = 3) according to manufacturer’s instructions (Abcam; ab100663 – VEGF human ELISA kit) using a Spark microplate reader /Tecan, Austria) at OD 450 nm.

### EV uptake and signaling in immune cells

To test uptake of EVs in PBMCs by flow cytometry and confocal microscopy and to question a hypothetic cell tropism, EVs were labeled with BODIPY™ FL C5-ceramide complexed to BSA (5 µg/ml; Thermo Fisher, USA) for 1 h at 4°C. To remove unbound dye, samples were diluted 1:10 with 0.9% NaCl solution containing 10mM HEPES and centrifuged at 4°C with 110,000 x g for 90 min in a Sorval WX80 ultracentrifuge with a TH-641 rotor at 25,400 rpm all (Thermo Fisher, USA). The pellet was re-suspended in the initial sample volume with 0.9% NaCl solution containing 10 mM HEPES. As a control for dye aggregates in the uptake experiment, dye in buffer alone was processed accordingly as a negative control sample. Particle concentration was determined thereafter by TRPS to adjust EV uptake number. PBMCs were cultured overnight in RPMI supplemented with 10% human AB-serum, 5 mM dipeptiven, 10 nM HEPES and 100 U/mL penicillin and 0.1 mg/mL streptomycin, in an Erlenmeyer flask at a density of 1 x 10^6^ cells/mL. After washing in PBS, unspecific binding was blocked using sheep serum and cells were stained with CD3-eF450 (2 µg/mL), CD19-APC (0.25 µg/mL), CD56-PE-Cy7 (0.5 µg/mL; all eBioscience, USA), CD14-APC-H7 (0.5 µg/mL; BD) and 7-AAD (1:100; eBioscience, USA). Washed cells were re-suspended in PBS containing 5% sheep serum and were sorted accordingly. Sort-purified cells were incubated with stained EVs for 24 h, fixed with 4% formalin and investigated for uptake with a laser scanning confocal microscope (Axio Observer Z1 attached to LSM700, Carl Zeiss).

To quantify the percentage and intensity of EV uptake, 300,000 PBMCs were incubated for 1 h, 24 h or 48 h with stained EVs or dye control sample in a ratio of EV to PBMC of 5,000:1. For flow cytometric quantification or EV uptake, PBMCs were washed with PBS, blocked with 2% sheep serum and stained with antibodies as indicated in the **Table S5**. Cells were analyzed with a 3-laser 10-color Gallios Flow cytometer (Beckman Coulter, USA).

Cell-type specific response to EV or soluble factor stimulation was investigated with sorted T cells and monocytes. Incubating equal sorted cell numbers for 15 min in a 10,000:1 ratio with TFF2 EVs or the corresponding volume of soluble factors was followed by cell lysis and analysis of the cell’s phosphor-kinome with the Human Phospho Kinase array Kit, ARY003C, (Human Phospho-RTK, R&D Systems, USA) according to manufacturer’s protocol.

### Skin grafting

Immune-deficient NOD.Cg-Prkdcscid Il2rgtm1WjI/SzJ mice (614NSG, Charles River) were used. Single-cell suspension transplants containing 6 x 10^6^ keratinocytes and 6 x 10^6^ fibroblasts diluted in 200 µL α-MEM/10% FBS (total volume 400 µL) were transplanted onto a full thickness back skin wound. Keratinocyte and fibroblasts isolation and cultivation, and the grafting procedure were performed as previously described^28^. In order to evaluate EV– induced angiogenesis, 200 µL of TFF2 PLX-EVs or equivalent volume of soluble factors from PLX cells obtained after separating TFF1s, cultivated in α-MEM/10% particle-depleted FBS, were added to the cell mixture before grafting. Skin biopsies were taken 14 days after initial grafting, fixed in 4% formaldehyde and prepared for histology.

### Histology of skin sections, image acquisition and quantification

For histochemistry and immune-histochemistry, paraffin-embedded skin samples were cut into 4 µm sections. Hematoxylin and eosin (HE) staining using Mayer’s Hemalaun (1.09249.2500, Merck) and Eosin Y (1.15935.0100, Merck) was done in a linear slide stainer (Leica ST4040). For Masson-Goldner trichrome staining, kits were used according to the manufacturer’s recommendations (12043, 14604, Morphisto). Automatic scanning of full slides in 40x magnification was done using the VS-120-L Olympus slide scanner 100-W system and processed using the Olympus VS-ASW-L100 program. Evaluation of the epidermal thickness and murine vessel quantification was done according to published work^28^, both measured in the Olympus VS-ASM-L100 program. Per group, four biological and at least three technical replicates were included.

### EV corona removal by ultracentrifugation and re-establishment of an EV corona

To deplete the EV protein corona, EV^TFF2^ preparations they were diluted 1:10 in NaCl 0.9% buffered with 10 mM HEPES and pelleted via ultracentrifugation at 4°C, 110,000 x g, for 90 min in a Sorval WX80 ultracentrifuge with a TH-641 rotor at 25,400 rpm all (Thermo Fisher, USA). The resulting pellet was re-suspended in the initial sample volume with NaCl 0.9% buffered with 10 mM HEPES. To re-establish a protein corona on the ‘naked’ EV^TUCF^ they were incubated for 1 h at 37°C in EGM-2 (Lonza, Basel, Switzerland) 4% of human serum albumin (Fresenius Kabi, Austria) and VEGF, IGF and EGF (Lonza, Basel, Switzerland) at dilutions indicated in the figure legend.

### Statistics

Statistical analysis of the results was performed using One-Way ANOVA analysis of variance with a confidence interval of 95% and corrected for multiple comparisons using the Holm Sidak algorithm in GraphPad Prism version 7.03. Proteomic results were analyzed using R. A p value of < 0.05 was defined as significant.

## Acknowledgements

The authors are grateful to Anna Hoog and the core facility flow cytometry at the PMU for excellent analysis support and to Sigrid Kahr from the University Hospital, Institute of Pathology in Salzburg, Austria for excellent technical assistance.

---

**Figure S1.**
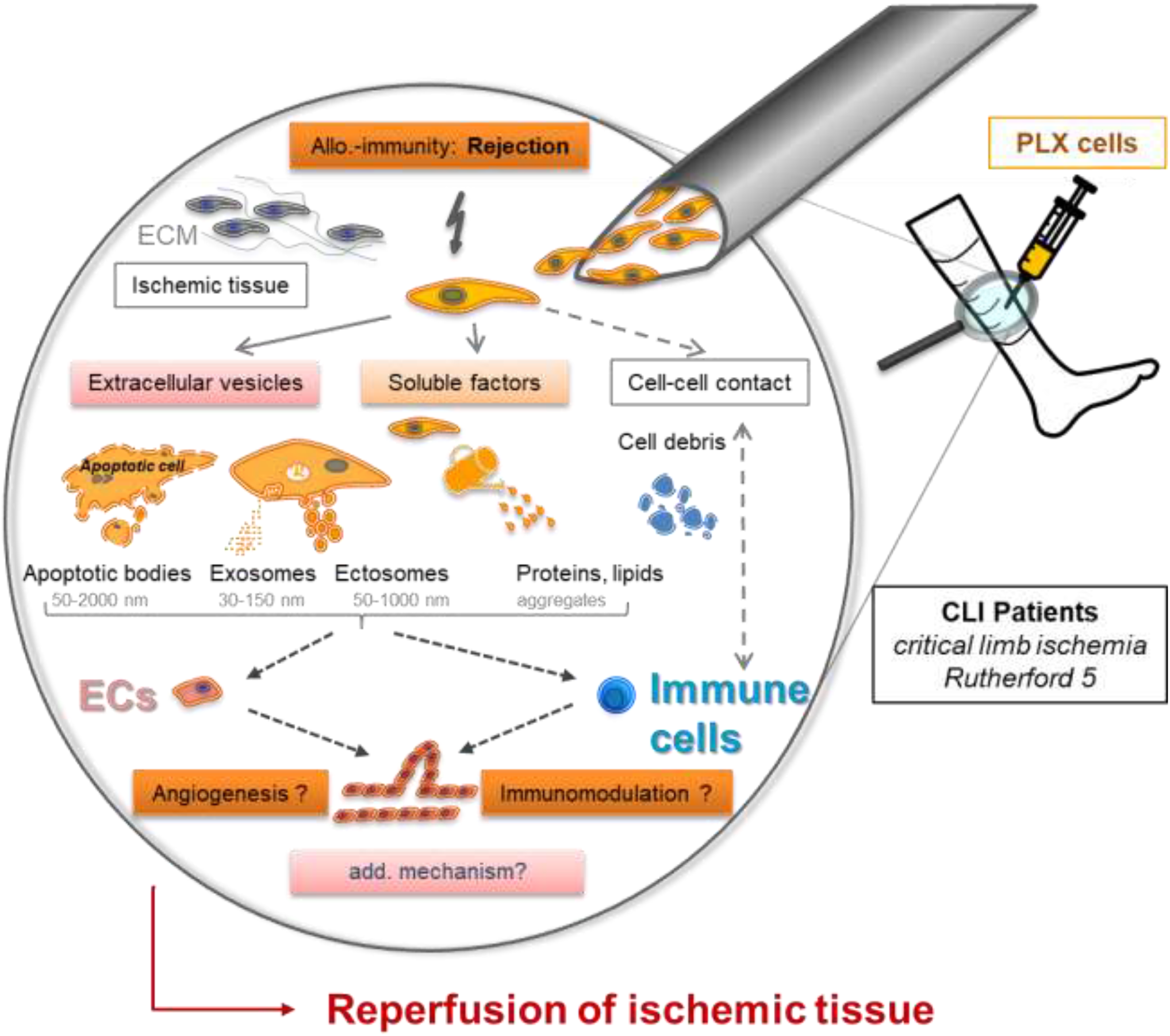
Hypothetic mode of action of allogeneic PLX stromal cells. Allogeneic placental-expanded (PLX) cells are considered to be rejected by the host immune system after local injection. The mode of action may include a temporary direct (cell-cell contact) or indirect stimulation of endogenous (endothelial or perivascular or interstitial) progenitor cells as well as immune response modulation by various types of secreted factors eventually resulting in reperfusion of the ischemic tissue and ideally wound healing. In this study, we focused on extracellular vesicles (EVs) and their protein cargo, compared to secreted soluble factors as potential mediators of PLX’s biologic activity.

**Figure S2:**
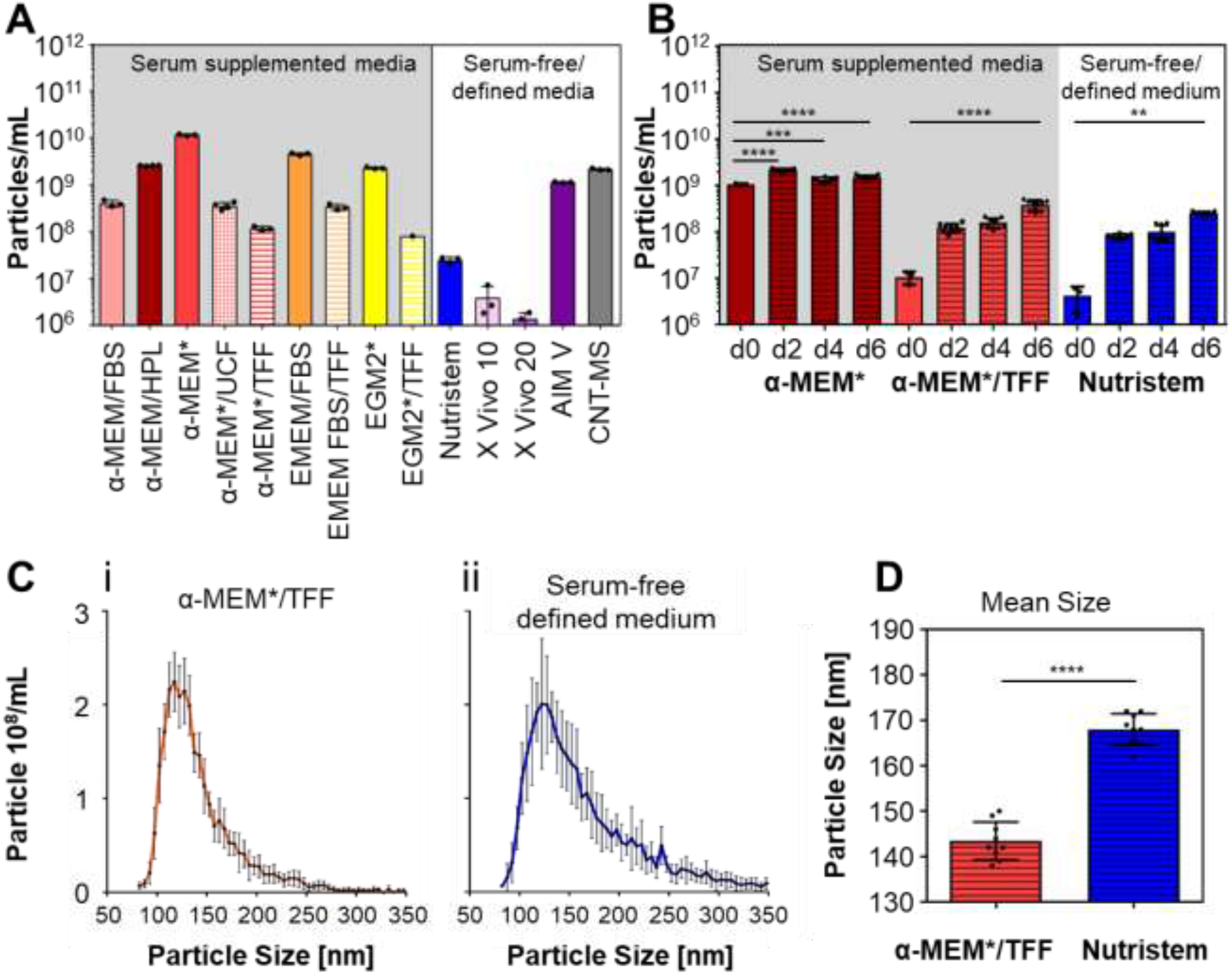
Physically defined media are required for cell-derived EV characterization. (**A**) Particle content of selected cell culture media with (left) or without serum (right) was measured by tunable resistive pulse sensing (TRPS). Media details are given in the methods section. (**B**) Comparison of the particle count in fresh and conditioned media after six-day placental stromal cell culture (PLX+). Particle concentrations were measured from three independent donors in four different media. (**C**) Time course of EV production in α-MEM*/TFF (*see* Fig. 1) was compared to serum free defined medium. TRPS analysis was performed for three independent donors in triplicates over six days. (**D**) TRPS measurement of PLX stromal cell-secreted EVs showed (**iii**) significantly different particle size distribution in (**ii**) defined serum-free vs. (**i**) platelet lysate-supplemented media (TFF pre-depleted). Triplicate measurements from one representative donor are shown (i, ii). Data from three donors were analyzed in triplicate (iii).

**Figure S3:**
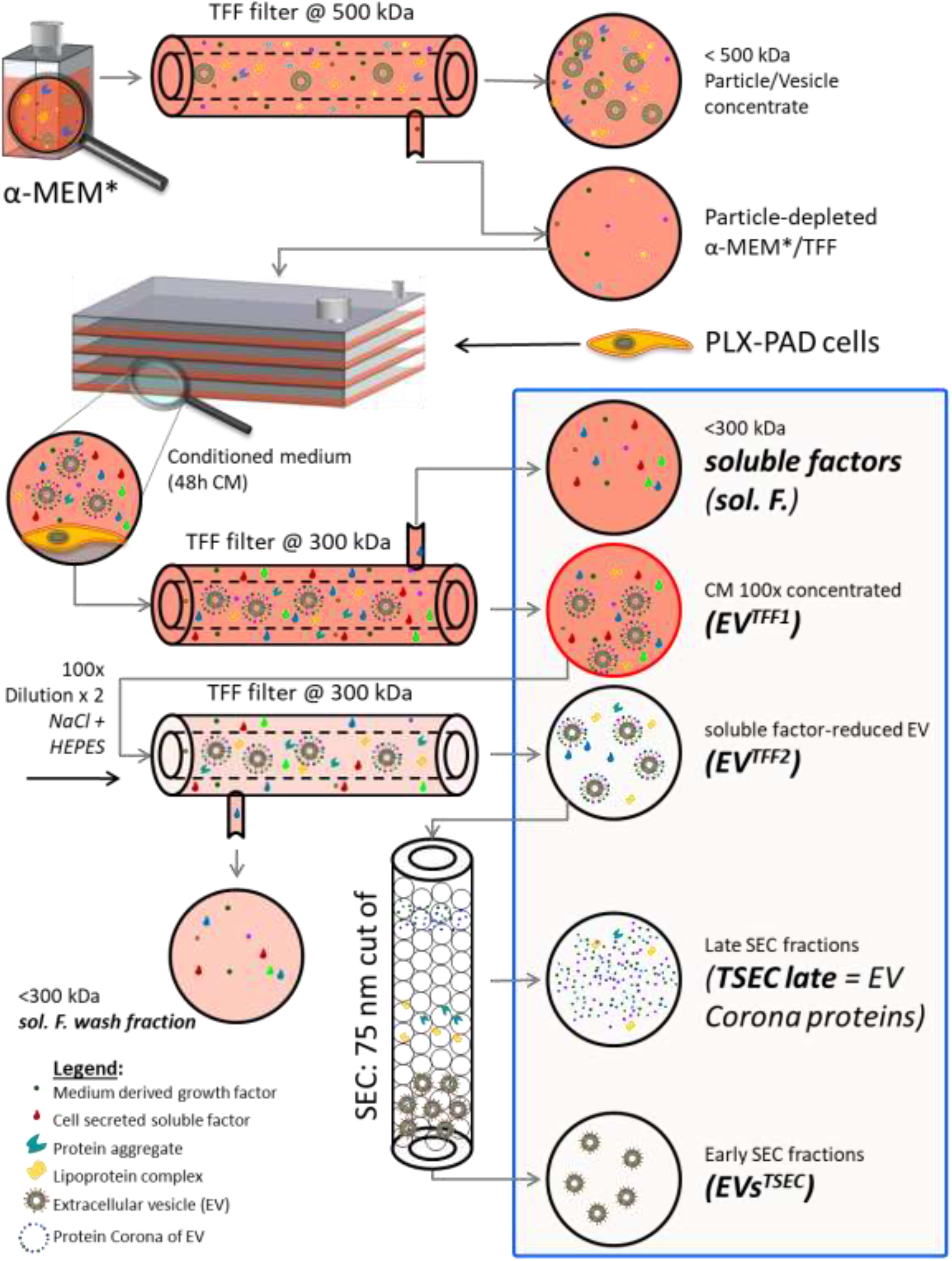
Schematic workflow of large-scale EV production from PLX cells. Cell culture medium α-MEM was supplemented with 10% pooled human platelet lysate (HPL) and fibrinogen-depleted (α-MEM*) before filtration through a 500 kDa cut-off membrane. 70% confluent PLX cells were cultured in particle-depleted α-MEM*/TFF for one to three 48-hour periods to obtain conditioned medium (CM). PLX-EVs were enriched 100-fold by TFF concentration (TFF1s) and separated from soluble factors (sol. F.). To remove remaining soluble factors/proteins, this crude concentrate was washed with twice the initial start volume to yield protein-TFF2 EVs separated from remote soluble factors. Size exclusion chromatography (SEC) was used to deplete EV-co-enriched soft corona proteins for selected experiments.

**Figure S4:**
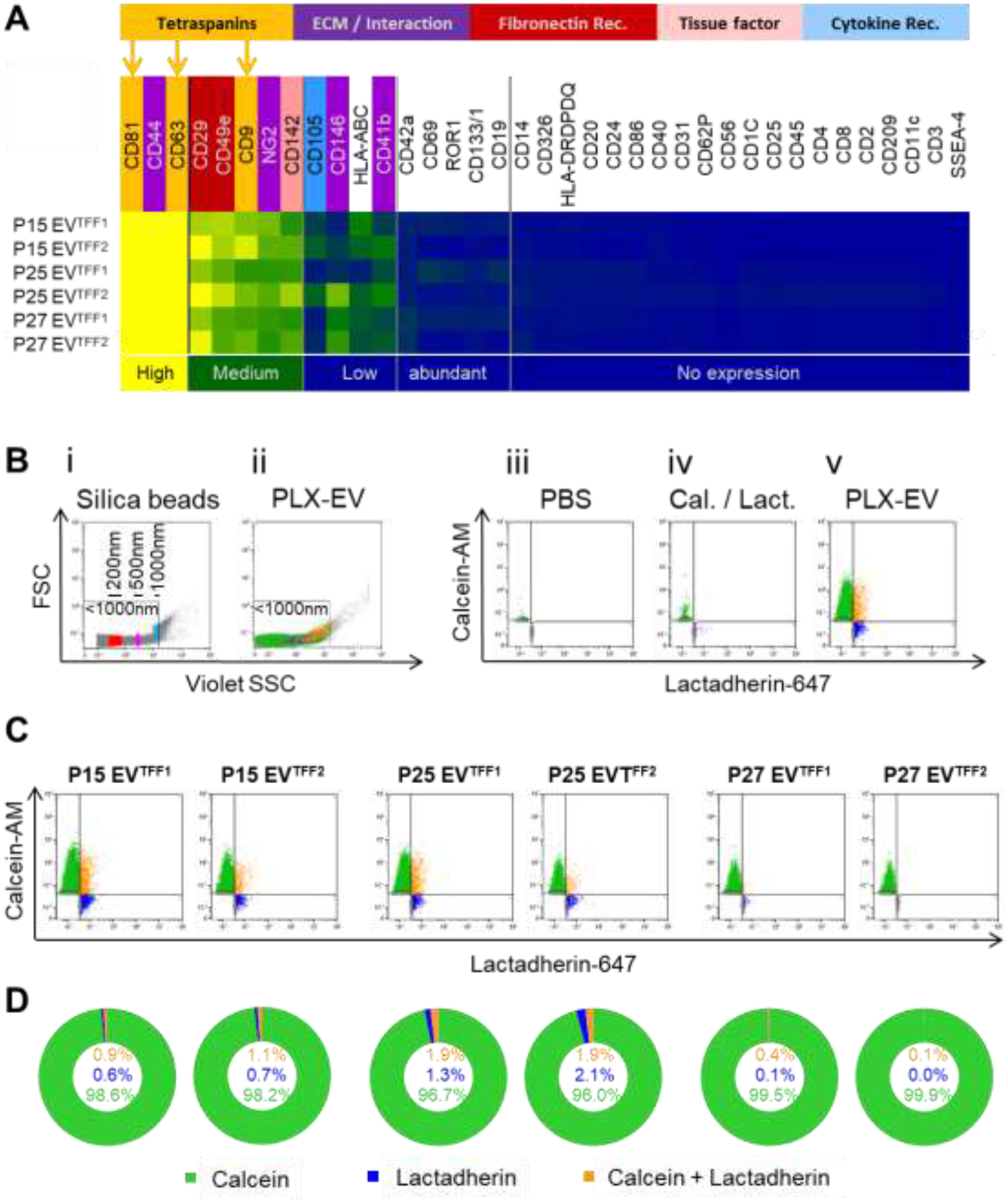
Antibody-based EV marker profiling indicated distinct address code and low abundance of apoptotic bodies. (**A**) Heat map showing mean surface marker expression of PLX-EVs from three individual donors (P15, P25, P27) as determined by bead-based multiplex flow cytometry (MACSPlex). (**B, i**) Fluorescent silica beads (Kisker) were used to adjust small particle resolution in flow cytometry and to set a size gate < 1,000 nm for EV detection. (**B, ii**) Dot plot showing size distribution of a representative PLX-EV sample. (**B, iii**) Dot plots showing fluorescence background of unstained PBS control, (**B, iv**) calcein-AM and lactadherin-Alexa Fluor 647 stained PBS control without EVs added (Cal. / Lact.) and (**B, v**) representative double-stained PLX-EV plot showing the distribution of calcein^+^ and lactadherin^+^ events using a dual fluorescence trigger. (**C**) Determination of calcein^+^ EVs and lactadherin^+^ presumably apoptotic bodies in crude vs. purified PLX-EV preparations. Dual fluorescence triggering-derived dot plots based on the gating strategy shown in (B) and color code as in (D) from three independent donor pairs of PLX-EV preparations from short-term PLX cultures of placenta lots P15, P25 and P27, respectively. (**D**) Pie charts depicting marker distribution of the positive events. Negative events were excluded from analysis because current technology does not permit precise discrimination between electronic noise and non-fluorescent unstained EVs or other undetermined non-particulate non-fluorescent signals notably in the size range of 100 nm and below ^44^.

**Figure S5:**
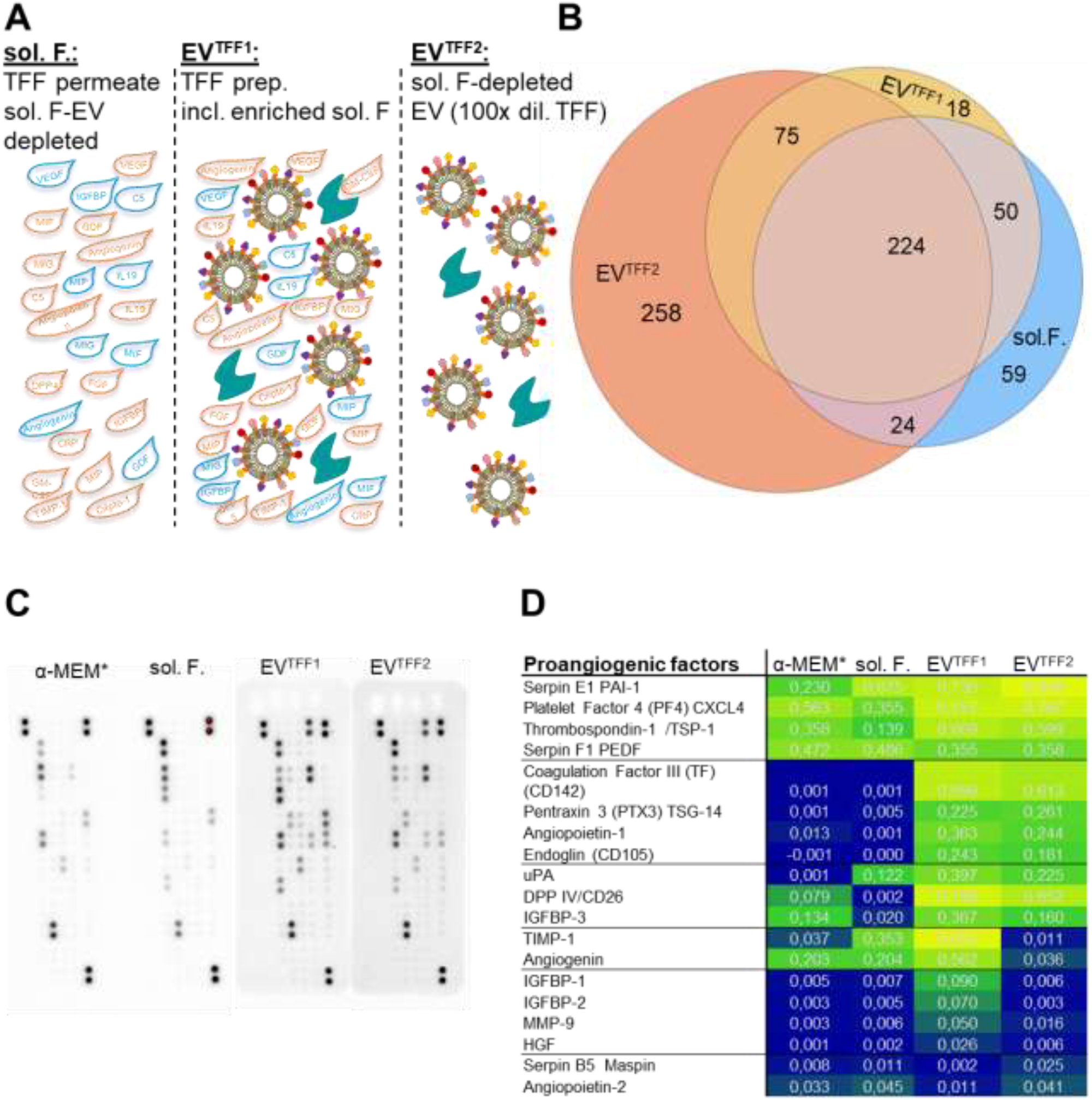
Proteomic composition of PLX secretome fractions. (**A**) Illustration of different fractions separated by TFF. A graphic symbol legend is shown in Figure 1. (**B**) Venn diagram showing the overlap and differences in proteins identified by label-free proteome analysis in the different secretome fractions (soluble factors, sol. F.; TFF1; purified EV, pur. EV) from three donors in two independent experiments. (**C**) Antibody array-based analysis of proangiogenic factors in the different secretome fractions (derived from one representative donor). (**D**) Heat map representation of corresponding quantitative analysis of PLX secretome fractions as indicated. Numbers represent relative luminescence units.

**Figure S6:**
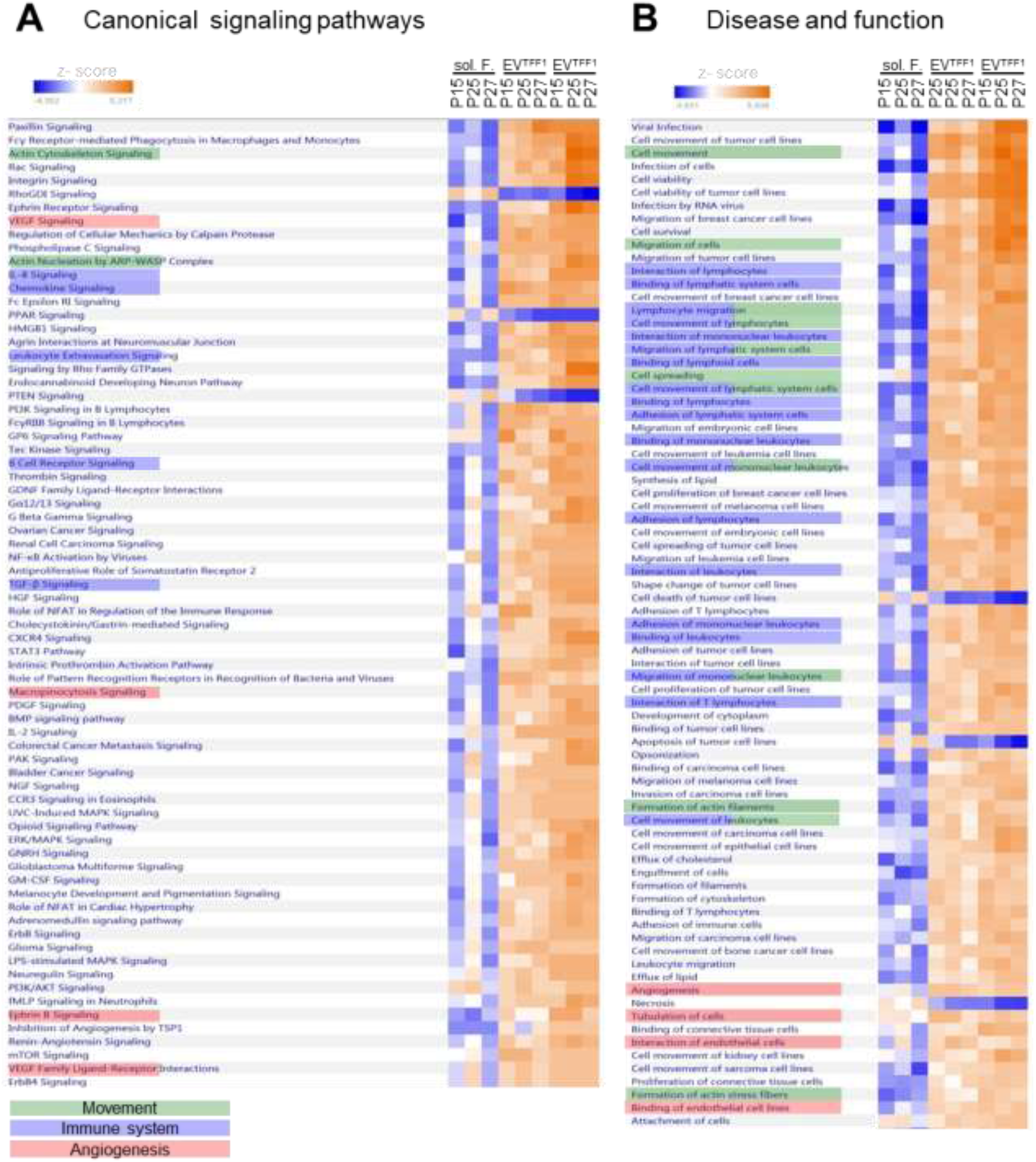
Ingenuity pathway (IPA) enrichment analysis of quantitative proteomics of the PLX secretome fractions. (**A**) Canonical signaling pathways differentially abundant in EVs versus soluble factors (sol.F.) are shown in a heatmap for three individual donors (P15, P25, P27) analyzed. Pathways related to movement, the immune system and angiogenesis are colored green, blue and red, respectively, as indicated. (**B**) Disease and function categories from the ingenuity database with the greatest differences in corresponding protein abundances between EVs and soluble factor fractions are shown in a heatmap using the same color code for highlighting pathways.

**Figure S7:**
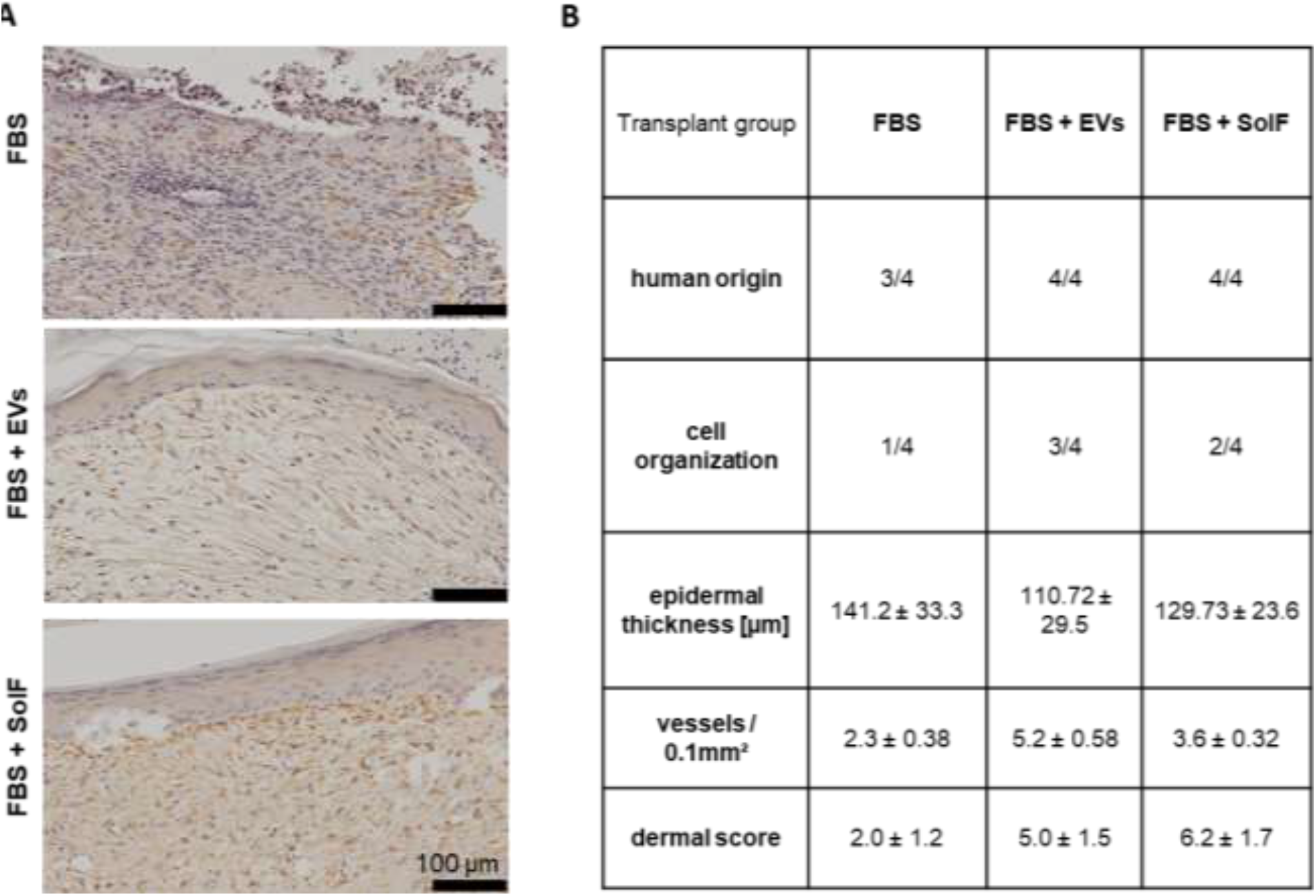
PLX-EVs promote skin regeneration in vivo. (**A**) Histology of human skin cell grafts on mice stained with anti-human vimentin antibody to verify human origin of the graft. Scale bar 100 µm. (**B**) Overview of important parameters to determine skin quality: human transplant establishment, proper organization of the different skin layers determined by scoring, epidermal thickness, vessel density/0.1mm² and overall dermal score^28^.

**Figure S8:**
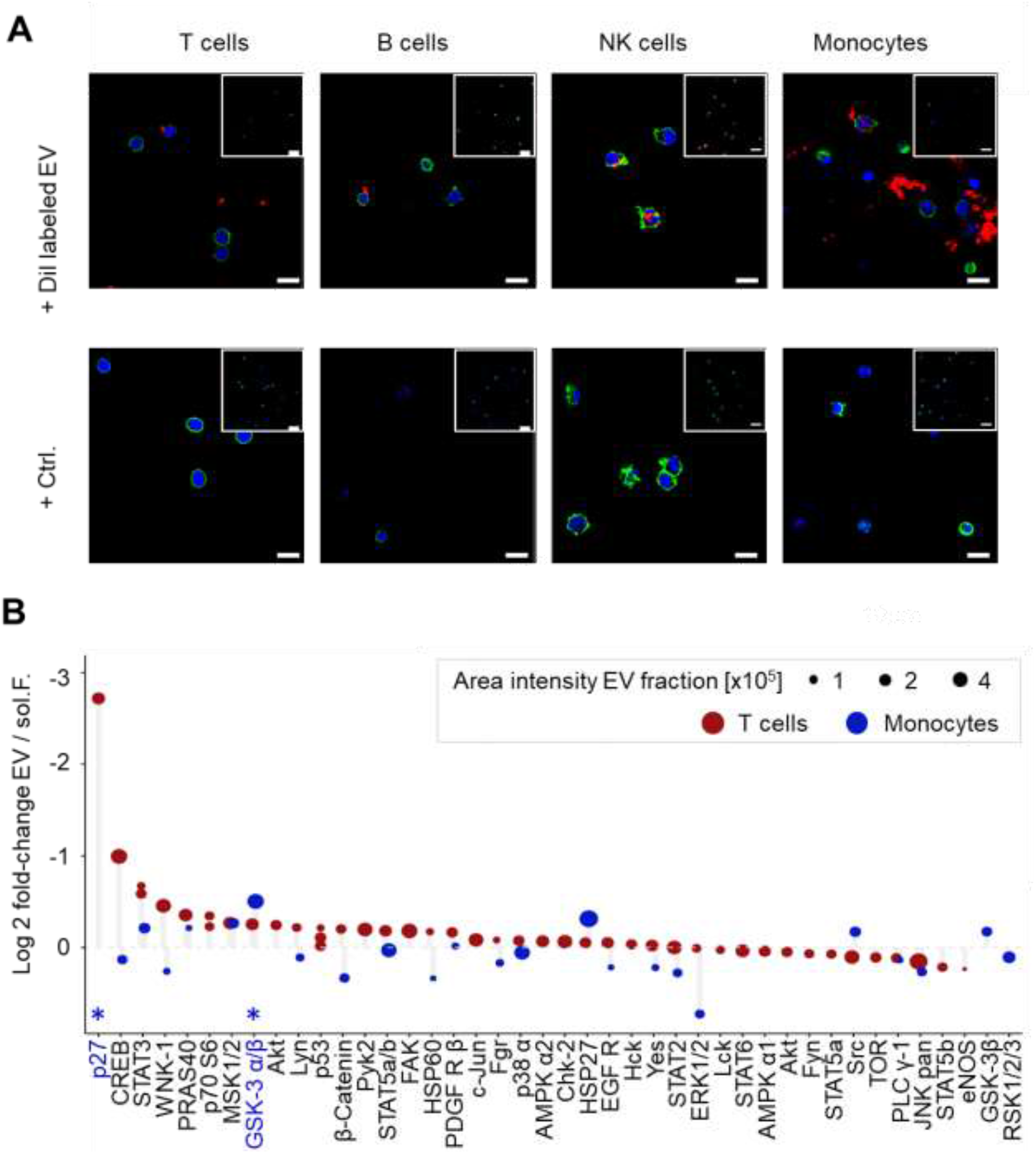
Immunomodulation and cell signaling by EVs: (**A)** Representative overview area after incubation of sorted T cells, B cells, NK cells and monocytes, stained with phalloidine (green) and DAPI ( blue) and incubated for 24 hours with Dil (red) labeled PLX EVs or control (Ctrl.) is shown. Scale bar 10µm. Insert represents overview with 40 µm scale-bar. (**B**) Sorted T cells and monocytes were treated with protein-TFF2 EVs for 15 minutes before cell lysis and kinome profiling as described in methods (n = 3). Significant changes for T cells are highlighted in blue text (p27 and GSK-3 α/β) as analyzed by pairwise T tests (*p < 0.05).

**Figure S9A:**
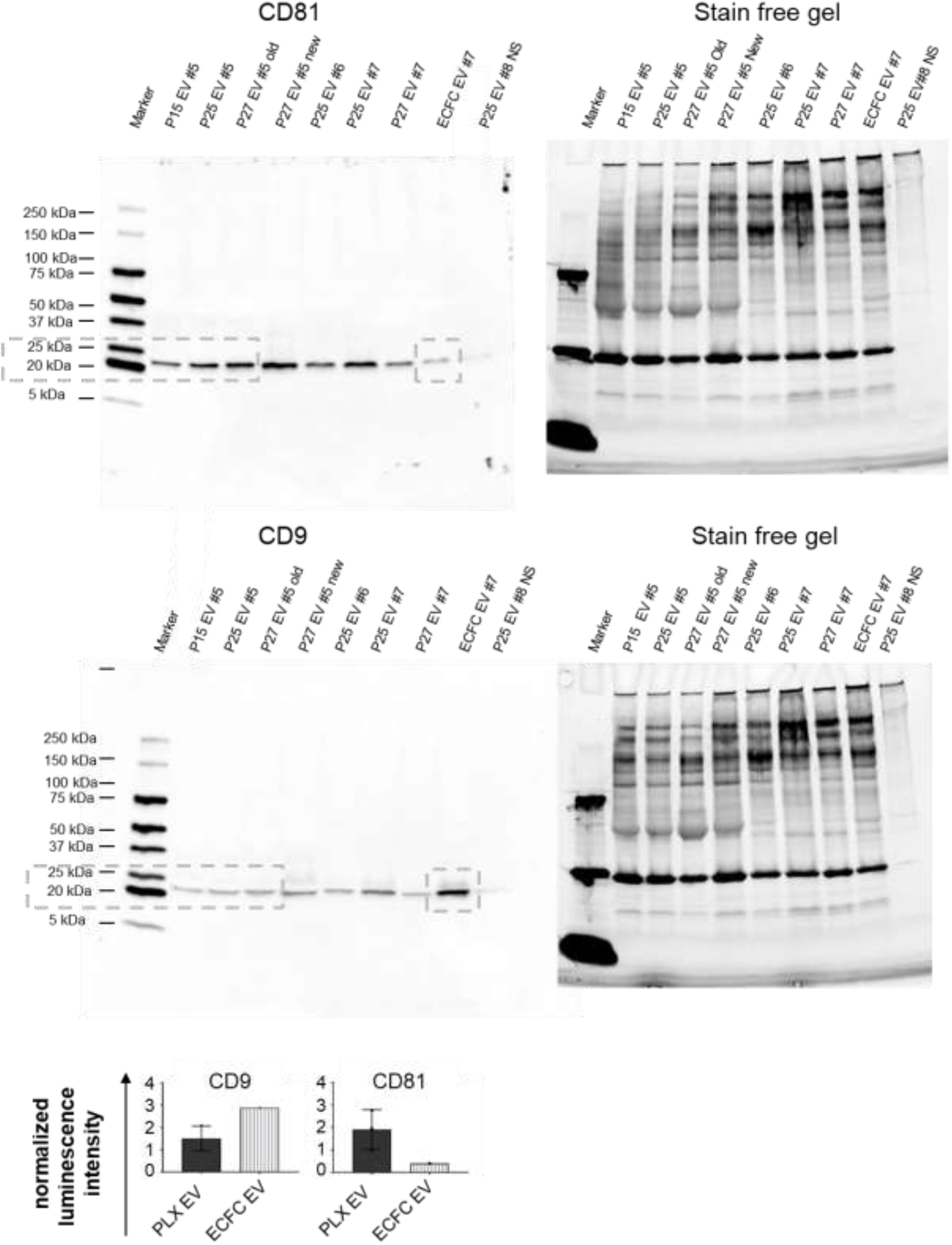
EV identity and purity determined by western blot: Whole membrane and gel of CD81 and CD9 immunoblots showing endothelial vs. stromal cell lineage-specific tetraspanin signature.

**Figure S9B:**
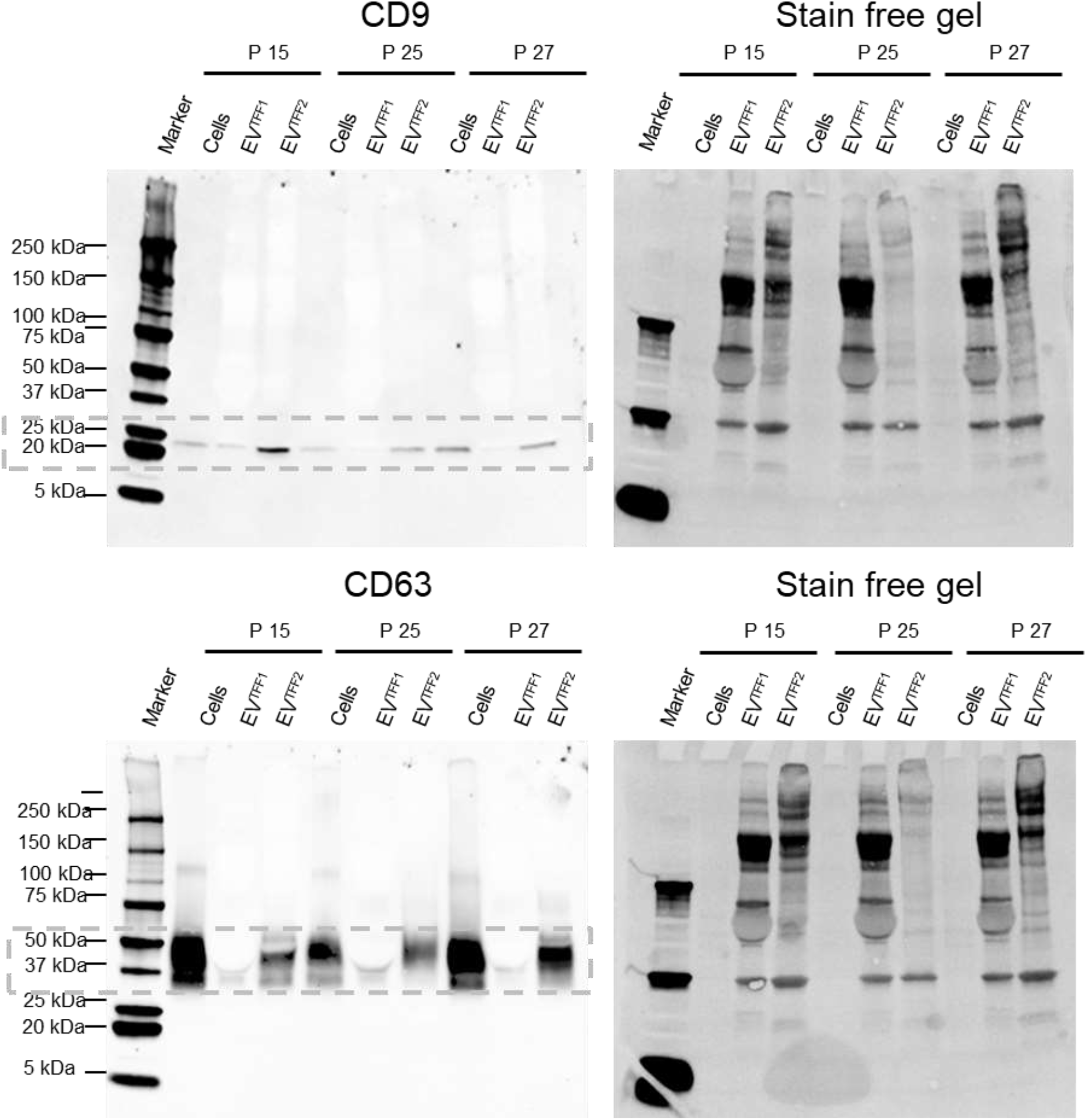
EV identity and purity determined by western blot: Whole membrane and gel of CD9 and CD63 immunoblots in Figure 1C.

**Figure S9C:**
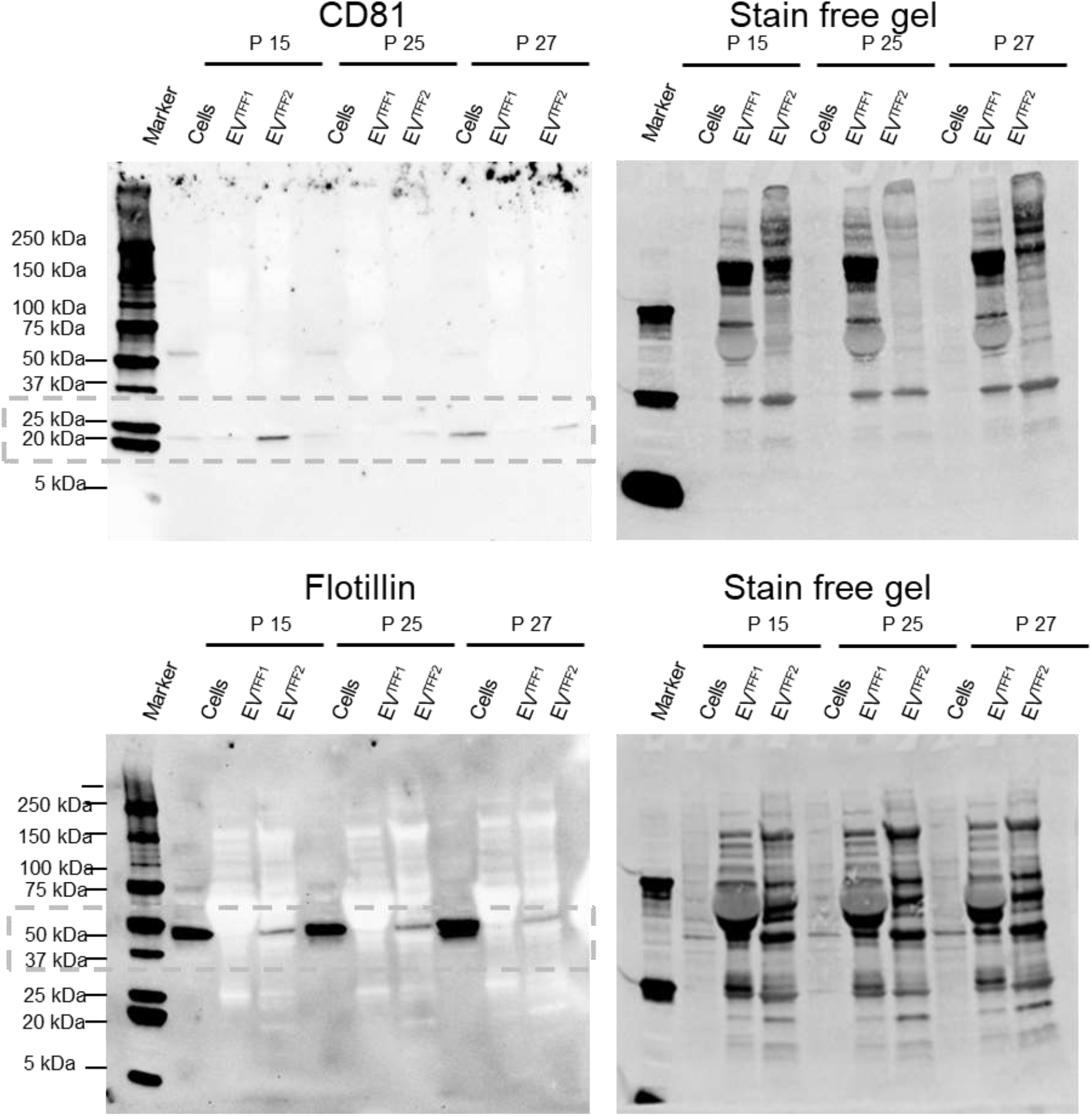
EV identity and purity determined by western blot: Whole membrane and gel of CD81 and flotillin immunoblots in Figure 1C.

**Figure S9D:**
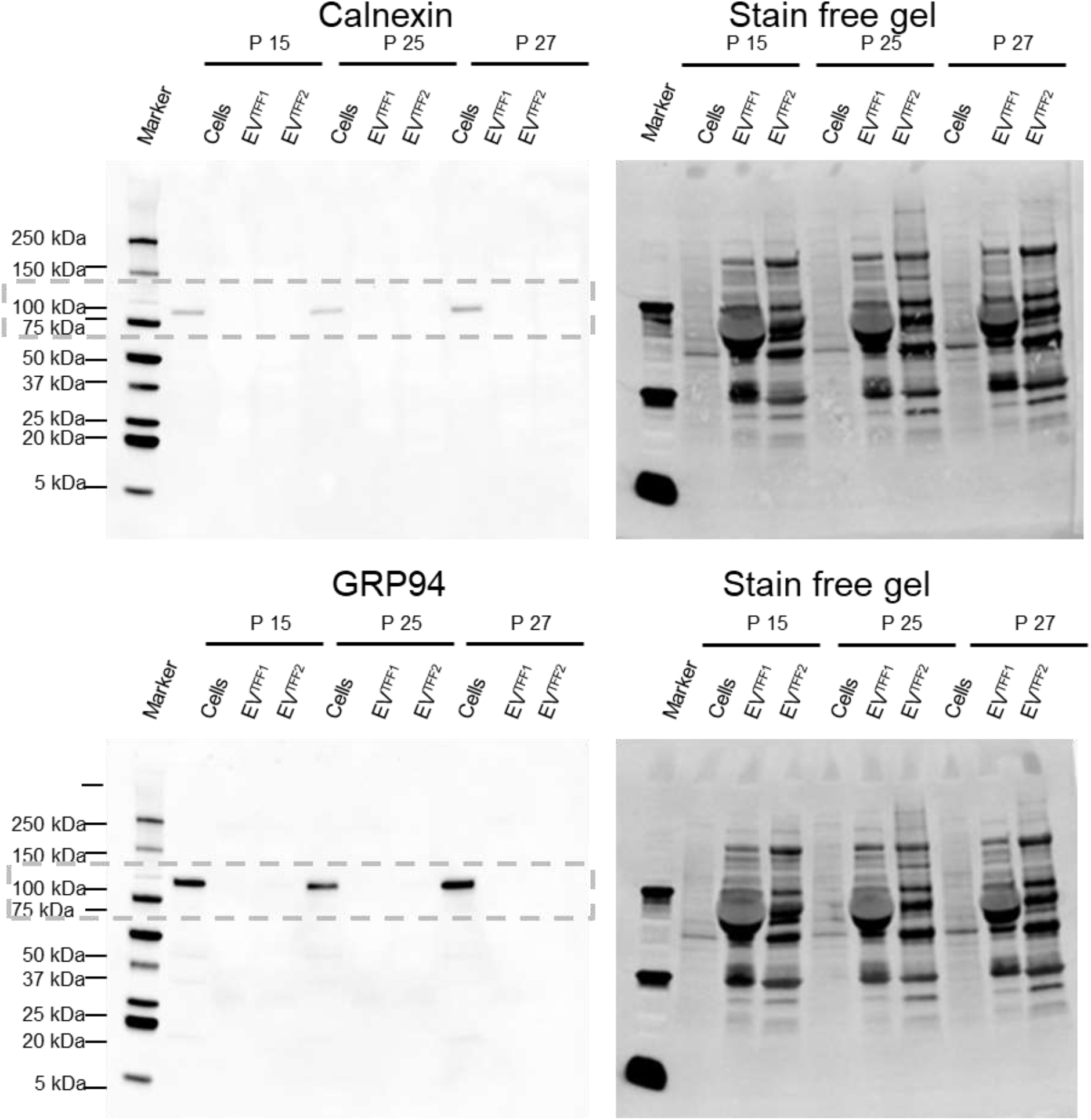
EV identity and purity determined by western blot: Whole membrane and gel of calnexin and GRP94 immunoblots in Figure 1C.

**Figure S9E:**
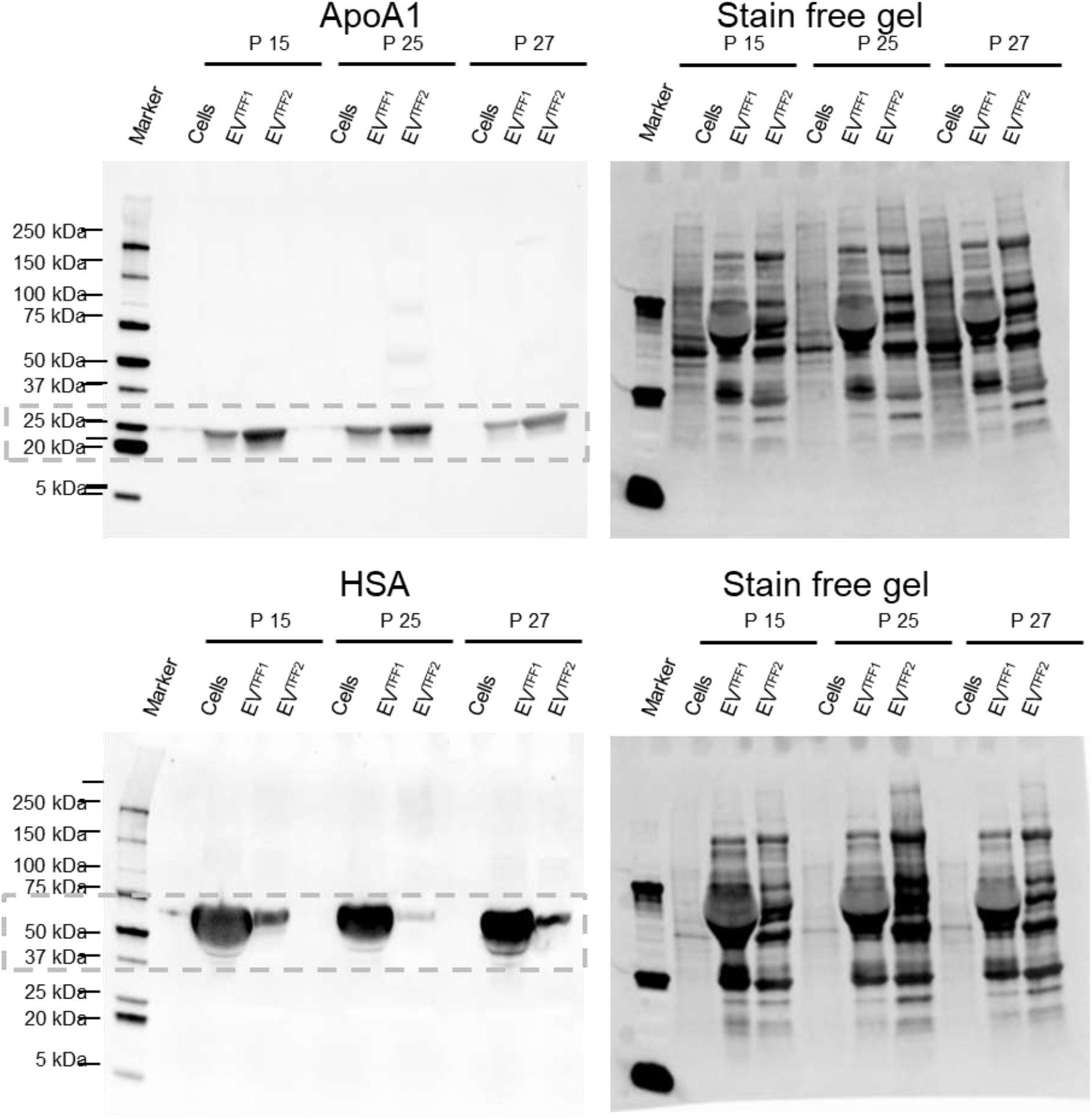
EV identity and purity determined by western blot: Whole membrane and gel of ApoA1 and HSA immunoblots in Figure 1C.

**Table S1:**
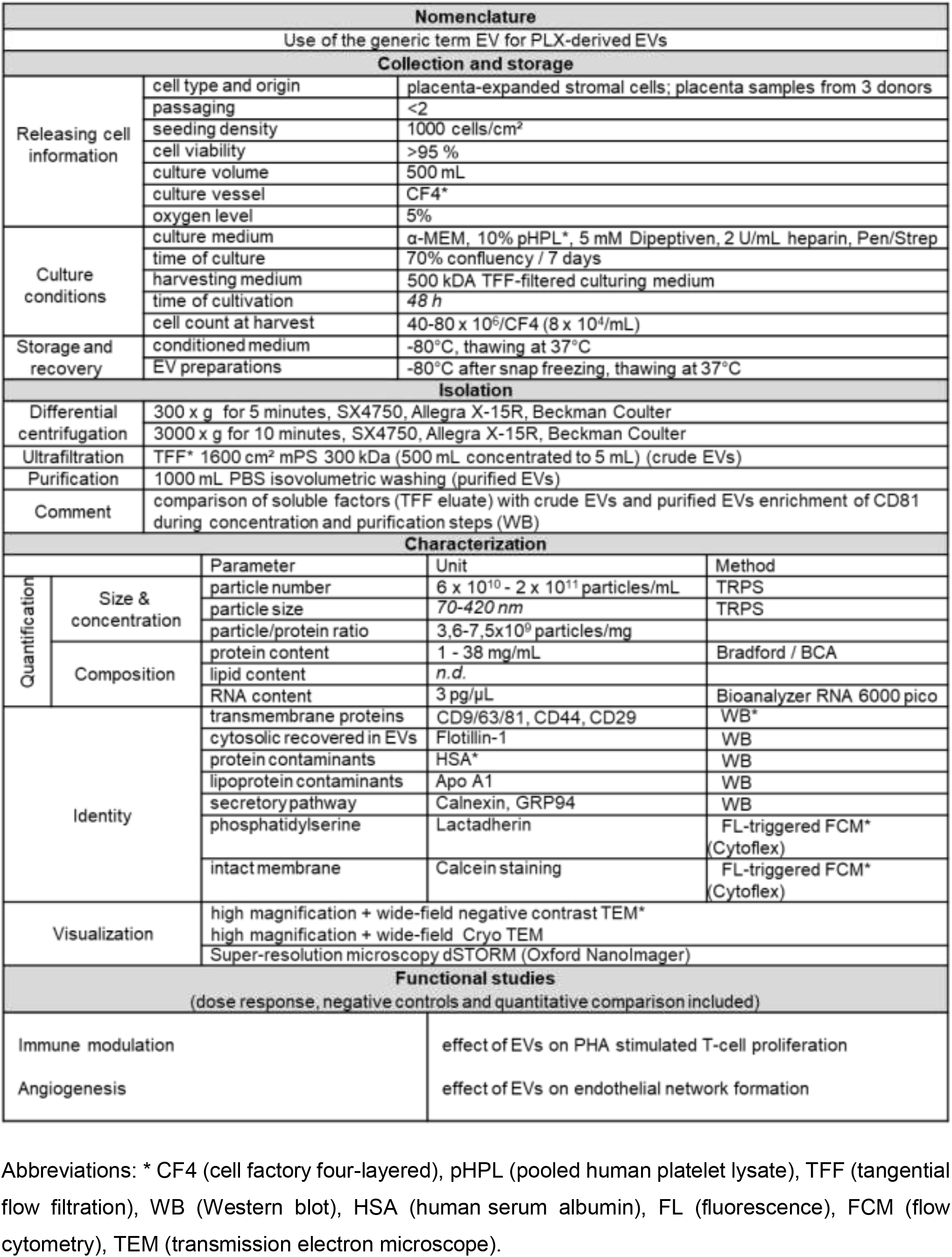
Addressing MISEV 2018 criteria for research with EV material in this study.

**Table S2:**
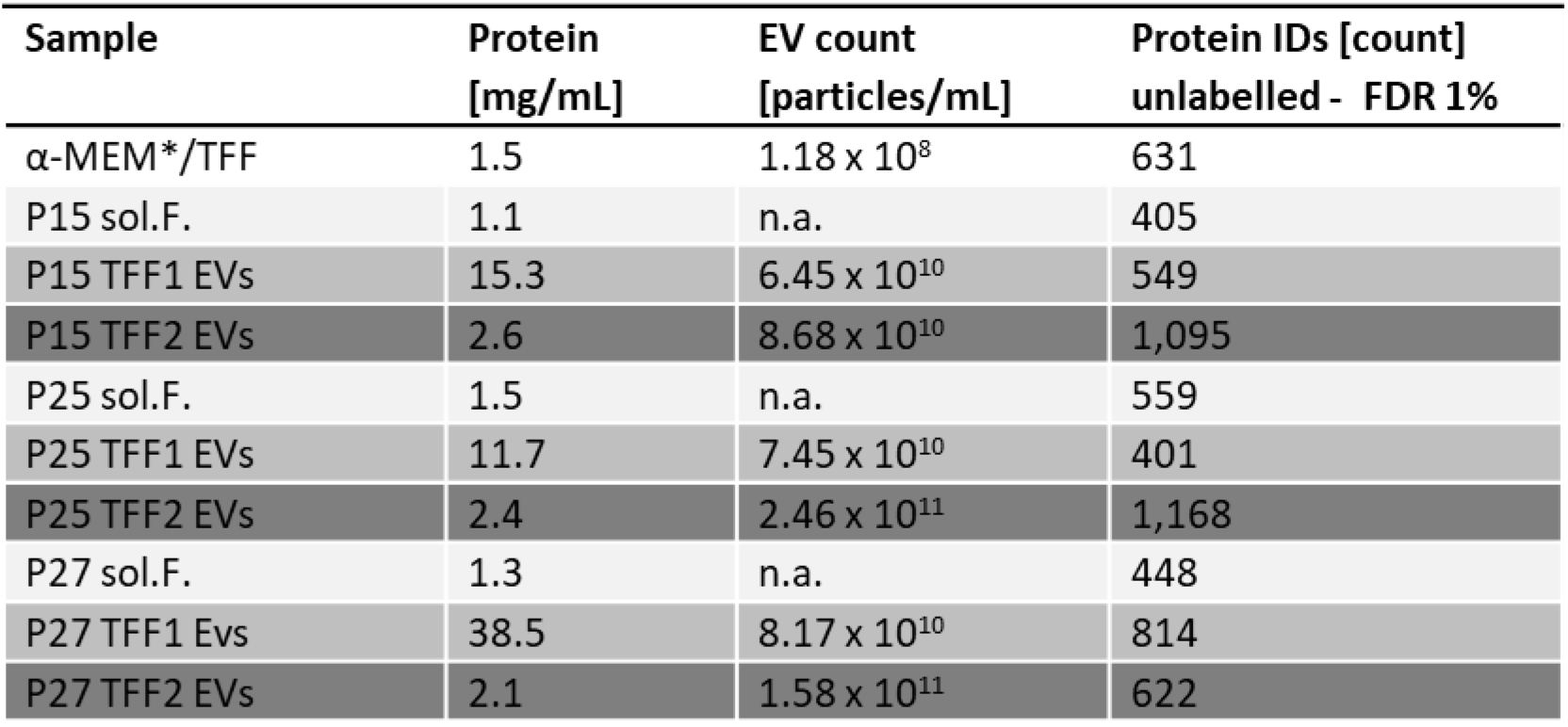
Sample details for proteomic analysis: Protein content as determined by BCA protein assay and particle count as measured by TRPS for samples used for proteomic analysis.

| Sample | Protein [mg/mL] | EV count [particles/mL] | Protein IDs [count] unlabelled - FDR 1% |
| --- | --- | --- | --- |
| α-MEM*/TFF | 1.5 | $1.18 \times 10^8$ | 631 |
| P15 sol.F. | 1.1 | n.a. | 405 |
| P15 TFF1 EVs | 15.3 | $6.45 \times 10^{10}$ | 549 |
| P15 TFF2 EVs | 2.6 | $8.68 \times 10^{10}$ | 1,095 |
| P25 sol.F. | 1.5 | n.a. | 559 |
| P25 TFF1 EVs | 11.7 | $7.45 \times 10^{10}$ | 401 |
| P25 TFF2 EVs | 2.4 | $2.46 \times 10^{11}$ | 1,168 |
| P27 sol.F. | 1.3 | n.a. | 448 |
| P27 TFF1 EVs | 38.5 | $8.17 \times 10^{10}$ | 814 |
| P27 TFF2 EVs | 2.1 | $1.58 \times 10^{11}$ | 622 |

**Table S3:**
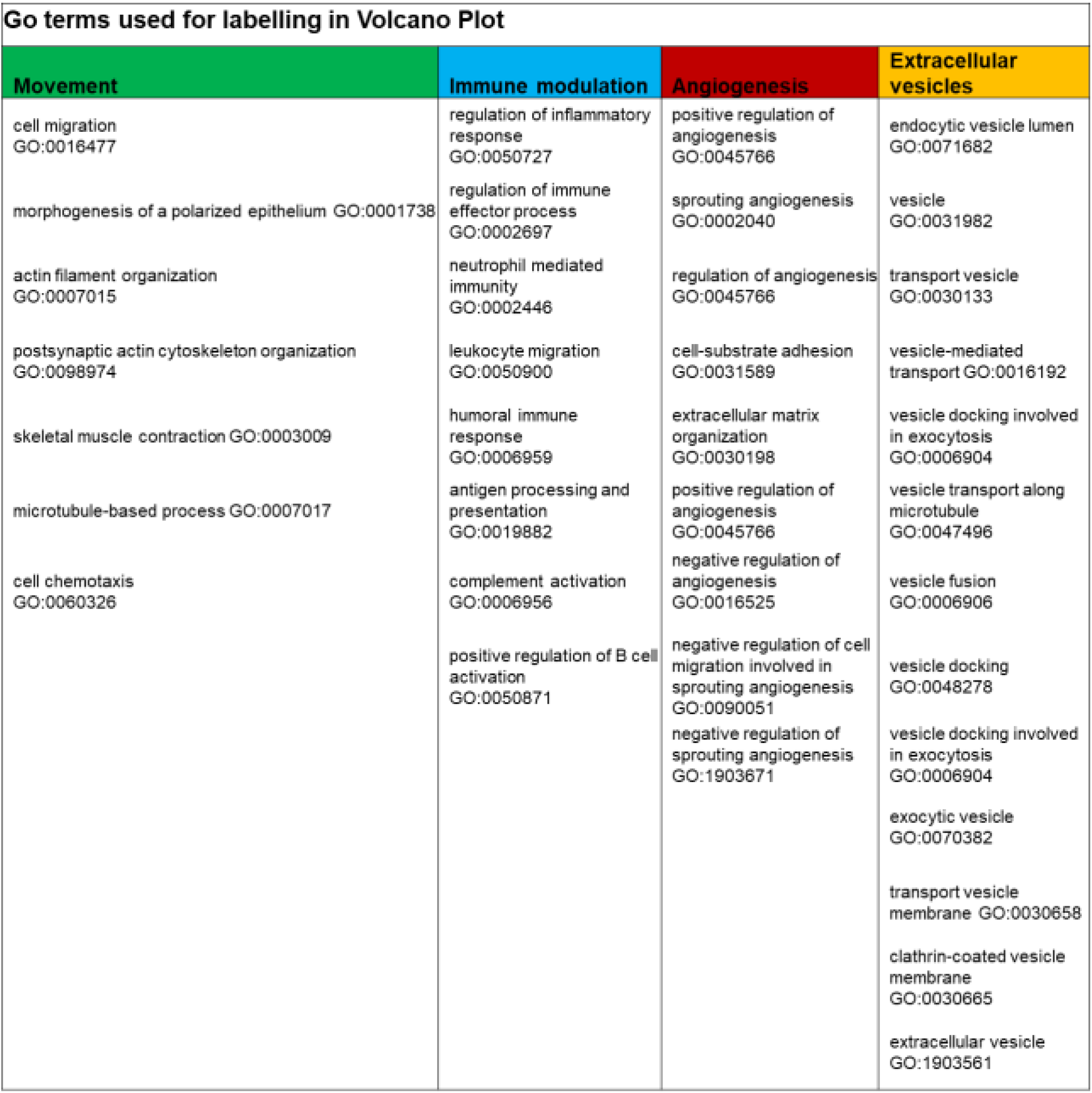
Quantitative proteomic characterization of PLX secretome fractions.

**Table S4:** Antibodies and dilutions used in western blots.

| Name | Supplier | Clone | Isotype | Concentration | WB dilution |
| --- | --- | --- | --- | --- | --- |
| **Apolipoprotein A1** | GeneTex | polyclonal | IgG | 0.66 mg/mL | 1:1320 |
| **Calnexin** | Cell Signaling | C5C9 | IgG | unknown | 1:1000 |
| **CD9** | Invitrogen | MM2/57IVA50 | IgG2 | 1 mg/mL | 1:50 |
| **CD63** | Thermo Fisher | TS63 | IgG1 | 0.5 mg/mL | 1:1000 |
| **CD81** | Bio-Rad | 1D6 | IgG1 | 1 mg/mL | 1:500 |
| **Flotillin 1** | BD | Flotillin-1 | IgG1 | 0.25 mg/mL | 1:1000 |
| **GRP94** | Bio-Rad | polyclonal | IgG | unknown | 1:2000 |
| **Human Serum Albumin** | Thermo Fisher | KT11 | IgG1 | 1 mg/mL | 1:1000 |

**Table S5:**
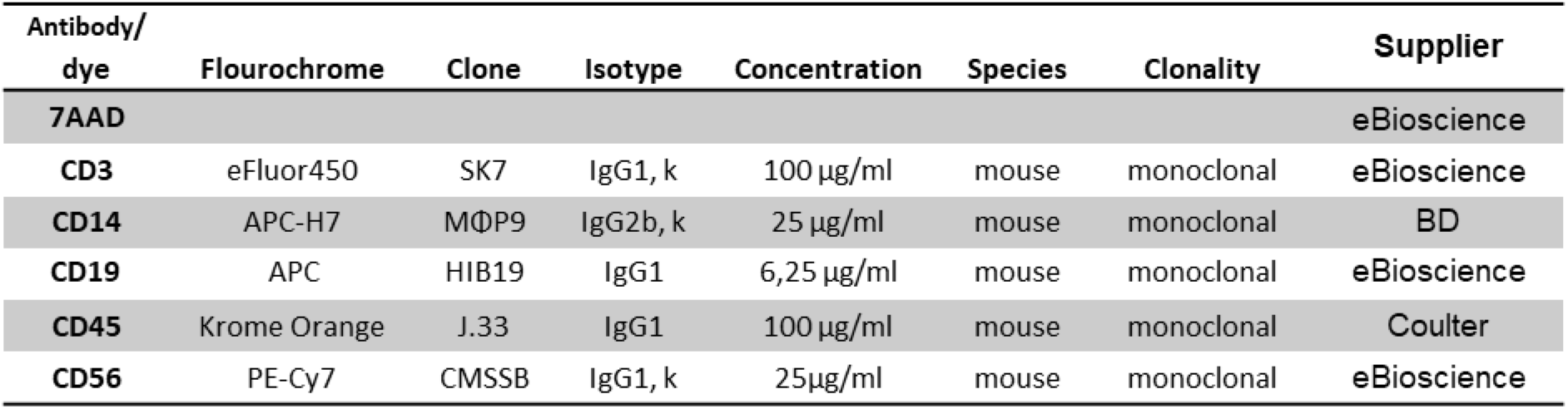
Antibodies and dilutions used for flow cytometry.

## References

1. Riazifar, M., Pone, E. J., Lotval, J. & Zhao, W. Stem Cell Extracellular Vesicles: Extended messages of regeneration. Annual Review of Pharmacology and Toxicology vol. 57 125–154 (2017).

2. Yáñez-Mó, M. et al. Biological properties of extracellular vesicles and their physiological functions. J. Extracell. Vesicles 4, 1–60 (2015).

3. Théry, C. et al. Minimal information for studies of extracellular vesicles 2018 (MISEV2018): a position statement of the International Society for Extracellular Vesicles and update of the MISEV2014 guidelines. J. Extracell. Vesicles 7, 1535750 (2018).

4. Van Niel, G., D’Angelo, G. & Raposo, G. Shedding light on the cell biology of extracellular vesicles. Nat. Rev. Mol. Cell Biol. 19, 213–228 (2018).

5. Neri, T. et al. CD18-mediated adhesion is required for the induction of a proinflammatory phenotype in lung epithelial cells by mononuclear cell-derived extracellular vesicles. Exp. Cell Res. 365, 78–84 (2018).

6. Cedervall, T. et al. Understanding the nanoparticle-protein corona using methods to quntify exchange rates and affinities of proteins for nanoparticles. Proc Natl Acad Sci USA 104, 2050–2055 (2007).

7. Ritz, S. et al. Protein Corona of Nanoparticles: Distinct Proteins Regulate the Cellular Uptake. Biomacromolecules 16, 1311–1321 (2015).

8. Tenzer, S. et al. Rapid formation of plasma protein corona critically affects nanoparticle pathophysiology. Nat. Nanotechnol. 8, 772–781 (2013).

9. Chan, K., Chao, S. & Kah, J. Exploiting Protein Corona around Gold Nanoparticles Conjugated to p53 Activating Peptides to Increase the Level of Stable p53 Proteins in Cells. Bioconjug. Chem. 30, 920–930 (2019).

10. Witwer, K. W. & Wolfram, J. Extracellular vesicles versus synthetic nanoparticles for drug delivery. Nat. Rev. Mater. 6, 103–106 (2021).

11. Song, P. et al. Global, regional, and national prevalence and risk factors for peripheral artery disease in 2015: an updated systematic review and analysis. Lancet Glob. Heal. 7, e1020–e1030 (2019).

12. Norgren, L. et al. PLX-PAD Cell Treatment of Critical Limb Ischaemia: Rationale and Design of the PACE Trial. Eur. J. Vasc. Endovasc. Surg. 57, 538–545 (2019).

13. Zahavi-Goldstein, E. et al. Placenta-derived PLX-PAD mesenchymal-like stromal cells are efficacious in rescuing blood flow in hind limb ischemia mouse model by a dose- and site-dependent mechanism of action. Cytotherapy 19, 1438–1446 (2017).

14. Pinzur, L. et al. Rescue from lethal acute radiation syndrome (ARS) with severe weight loss by secretome of intramuscularly injected human placental stromal cells. J. Cachexia. Sarcopenia Muscle 9, 1079–1092 (2018).

15. Winkler, T. et al. Immunomodulatory placental-expanded, mesenchymal stromal cells improve muscle function following hip arthroplasty. J. Cachexia. Sarcopenia Muscle 9, 880–897 (2018).

16. Qazi, T. H. et al. Cell therapy to improve regeneration of skeletal muscle injuries. J. Cachexia. Sarcopenia Muscle 10, 501–516 (2019).

17. Ankrum, J. A., Ong, J. F. & Karp, J. M. Mesenchymal stem cells: immune evasive, not immune privileged. Nat. Biotechnol. 32, 252–260 (2014).

18. Lener, T. et al. Applying extracellular vesicles based therapeutics in clinical trials - An ISEV position paper. J. Extracell. Vesicles 4, (2015).

19. Herrmann, I. K., Wood, M. J. A. & Fuhrmann, G. Extracellular vesicles as a next-generation drug delivery platform. Nat. Nanotechnol. 16, 748–759 (2021).

20. Schallmoser, K. & Strunk, D. Preparation of pooled human platelet lysate (pHPL) as an efficient supplement for animal serum-free human stem cell cultures. J. Vis. Exp. (2009) doi:10.3791/1523.

21. Colombo, M., Raposo, G. & Théry, C. Biogenesis, secretion, and intercellular interactions of exosomes and other extracellular vesicles. Annual review of cell and developmental biology vol. 30 255–289 (2014).

22. Laner-Plamberger, S. et al. Mechanical fibrinogen-depletion supports heparin-free mesenchymal stem cell propagation in human platelet lysate. J. Transl. Med. 13, 354 (2015).

23. Shelke, G. V., Lässer, C., Gho, Y. S. & Lötvall, J. Importance of exosome depletion protocols to eliminate functional and RNA-containing extracellular vesicles from fetal bovine serum. J. Extracell. Vesicles 3, (2014).

24. Pachler, K. et al. An In Vitro Potency Assay for Monitoring the Immunomodulatory Potential of Stromal Cell-Derived Extracellular Vesicles. International Journal of Molecular Sciences vol. 18 (2017).

25. Wiklander, O. P. B. B. et al. Systematic methodological evaluation of a multiplex bead-based flow cytometry assay for detection of extracellular vesicle surface signatures. Front. Immunol. 9, 1326 (2018).

26. Crescitelli, R. et al. Distinct RNA profiles in subpopulations of extracellular vesicles: Apoptotic bodies, microvesicles and exosomes. J. Extracell. Vesicles 2, 20677 (2013).

27. Wang, C. K., Nelson, C. F., Brinkman, A. M., Miller, A. C. & Hoeffler, W. K. Spontaneous Cell Sorting of Fibroblasts and Keratinocytes Creates an Organotypic Human Skin Equivalent. J. Invest. Dermatol. 114, 674–680 (2000).

28. Ebner-Peking, P. et al. Self-assembly of differentiated progenitor cells facilitates spheroid human skin organoid formation and planar skin regeneration. Theranostics 11, 8430–8447 (2021).

29. Kokkinopoulou, M., Simon, J., Landfester, K., Mailänder, V. & Lieberwirth, I. Visualization of the protein corona: towards a biomolecular understanding of nanoparticle-cell-interactions. Nanoscale 9, 8858–8870 (2017).

30. Kalluri, R. & LeBleu, V. S. The biology, function, and biomedical applications of exosomes. Science (80-.). 367, eaau6977 (2020).

31. Hadjidemetriou, M. & Kostarelos, K. Evolution of the nanoparticle corona. Nat. Nanotechnol. 12, 288–290 (2017).

32. Reinisch, A. et al. Humanized large-scale expanded endothelial colony – forming cells function in vitro and in vivo. Blood 113, 6716–6725 (2009).

33. Poupardin, R., Wolf, M. & Strunk, D. Adherence to minimal experimental requirements for defining extracellular vesicles and their functions. Advanced Drug Delivery Reviews vol. 176 (2021).

34. Dawson, K. A. & Yan, Y. Current understanding of biological identity at the nanoscale and future prospects. Nat. Nanotechnol. 16, 229–242 (2021).

35. Burnouf, T., Strunk, D., Koh, M. B. C. & Schallmoser, K. Human platelet lysate: Replacing fetal bovine serum as a gold standard for human cell propagation? Biomaterials vol. 76 371–387 (2016).

36. Reinisch, A. & Strunk, D. Isolation and animal serum free expansion of human umbilical cord derived mesenchymal stromal cells (MSCs) and endothelial colony forming progenitor cells (ECFCs). J. Vis. Exp. 1525 (2009) doi:10.3791/1525.

37. Schallmoser, K. et al. Human platelet lysate can replace fetal bovine serum for clinical-scale expansion of functional mesenchymal stromal cells. Transfusion 47, 1436–1446 (2007).

38. Bartmann, C. et al. Two steps to functional mesenchymal stromal cells for clinical application. Transfusion 47, 1426–1435 (2007).

39. Koliha, N. et al. A novel multiplex bead-based platform highlights the diversity of extracellular vesicles. J. Extracell. Vesicles 5, 1–15 (2016).

40. Mastronarde, D. N. Automated electron microscope tomography using robust prediction of specimen movements. J. Struct. Biol. 152, 36–51 (2005).

41. Cox, J. & Mann, M. MaxQuant enables high peptide identification rates, individualized p.p.b.-range mass accuracies and proteome-wide protein quantification. Nat. Biotechnol. 26, 1367–1372 (2008).

42. The UniProt Consortium. UniProt: the universal protein knowledgebase. Nucleic Acids Res. 45, D158–D169 (2017).

43. Ketterl, N. et al. A robust potency assay highlights significant donor variation of human mesenchymal stem/progenitor cell immune modulatory capacity and extended radio-resistance. Stem Cell Res. Ther. 6, 1–11 (2015).

44. Görgens, A. et al. Optimisation of imaging flow cytometry for the analysis of single extracellular vesicles by using fluorescence-tagged vesicles as biological reference material. J. Extracell. Vesicles 8, 1587567 (2019).

